# High exposure variance enables candidate biomarker detection in a small EWAS of methylmercury-exposed Peruvian adults

**DOI:** 10.1101/2022.07.05.498896

**Authors:** Caren Weinhouse, Luiza Perez, Ian Ryde, Jaclyn M. Goodrich, J. Jaime Miranda, Heileen Hsu-Kim, Susan K. Murphy, Joel N. Meyer, William K. Pan

**Affiliations:** Oregon Institute of Occupational Health Sciences, Oregon Health & Science University, Portland, Oregon; Duke Global Health Institute, Duke University, Durham, North Carolina; Environmental Science and Policy, Nicholas School of the Environment, Duke University, Durham, North Carolina; Department of Environmental Health Sciences, University of Michigan School of Public Health, Ann Arbor, Michigan; Sydney School of Public Health, Sydney, Australia; CRONICAS Center of Excellence in Chronic Diseases, Universidad Peruana Cayetano Heredia, Lima, Peru; Department of Civil and Environmental Engineering, Pratt School of Engineering, Duke University, Durham, North Carolina; Division of Reproductive Sciences, Department of Obstetrics and Gynecology, Duke University Medical Center, Durham, North Carolina

**Keywords:** DNA methylation, Epigenetics, Methylmercury, Mitochondrial DNA, Epigenetic age

## Abstract

I.

**Background:** Epigenome-wide association studies (EWAS) are a highly promising approach that can inform precision environmental health. However, current EWAS are underpowered for biomarker detection and increasing sample sizes will require substantial resources. Therefore, alternative approaches for identifying candidate biomarkers through EWAS are critical for moving the field forward.

**Objectives:** To provide proof-of-principle that maximizing exposure variance in EWAS by selecting participants from disproportionately exposed global populations enables effective candidate biomarker detection, even in small sample sizes.

**Methods:** We profiled genome-wide DNA methylation using Illumina Infinium MethylationEPIC BeadChip in whole blood from N=32 individuals from Madre de Dios, Peru with high methylmercury (MeHg) exposure due to artisanal and small-scale gold mining. We compared DNA methylation in N=16 individuals with high (>10 μg/g) vs. N=16 individuals with low (<1 μg/g) total hair mercury (a proxy for methylmercury exposure), matched on age and sex.

**Results:** We identified nine differentially methylated CpG sites (FDR<0.05), including several with known links to MeHg toxicity. The most significantly different CpG site was in an intronic enhancer of the *SLC5A7* gene, which encodes the L-type amino acid transporter 1 (LAT1) that facilitates MeHg transport into protein-rich tissue, including muscle and brain. Our Gene Ontology and transcription factor motif enrichment analyses identified differential methylation of genes involved in several outcomes with established links to MeHg, including immune response, neurotoxicity, and type 2 diabetes (T2D) risk. Last, we identified candidate epigenetic biomarkers of PUFA-mediated protection against MeHg toxicity.

**Discussion:** Here, we show that a small EWAS on samples with high MeHg exposure variance can detect candidate differentially methylated CpGs and pathways of interest relevant to MeHg biology. Similar EWAS in global populations with known high exposure variance can be leveraged to develop targeted, custom sequencing panels and microarrays limited to replicated, validated biomarkers of a given exposure.

## II. Introduction

Precision environmental health describes an individualized approach to public health protection that accounts for differences in individuals’ comprehensive exposures, or “exposomes”, and their biological responses to those exposures^1,2^. An important component of this approach is the development of biomarkers of exposure and effect that reflect individuals’ biological responses to chemical exposures^1,3^. One promising category of biomarkers is epigenetic biomarkers, primarily regions of DNA methylation of cytosines within cytosine-guanine dinucleotides (CpG sites) that reflect current or past gene expression responses to pollutants or nutrients^1,4,5^. Ideal epigenetic biomarkers are specific to particular chemical or dietary exposures and reliably report on biological responses that reflect known mechanisms of toxicity or protection^1,4^. Although exciting in theory, in practice, descriptive discovery experiments comparing people with high vs. low levels of environmental exposures have identified only a small number of candidate epigenetic biomarkers, likely due to insufficient statistical power to detect true differences between populations^5^. This limitation can be overcome with larger sample sizes; current epigenome-wide association studies (EWAS) for DNA methylation markers average sample sizes in the hundreds or low thousands^5^. For comparison, genome-wide association studies (GWAS) average sample sizes in the tens or hundreds of thousands^5^. This approach will improve statistical power but it will require substantial time and resources to implement. Therefore, it is important to consider alternative approaches for detecting candidate DNA methylation biomarkers in EWAS studies.

In this study, we provide proof-of-principle that maximizing exposure variance, rather than increasing sample size, is a potential alternative approach for candidate biomarker detection in EWAS studies. We conducted a discovery EWAS using the Illumina Infinium MethylationEPIC BeadChip in age- and sex-matched Peruvian adults with high variance in methylmercury (MeHg) exposure due to nearby artisanal and small-scale gold mining^6^ (**Table 1**). We show that a very small EWAS cohort (N=16 participants with high exposure vs. N=16 with low exposure) (**Table 1**) enables detection of an equivalent number of candidate DNA methylation biomarkers (N=9 differentially methylated CpG sites) (**Table 2**), as compared to existing, larger EWAS studies (N=1-20 differentially methylated CpG sites)^5^. In addition, our Gene Ontology and transcription factor motif enrichment analyses yielded substantial insights into biological responses to MeHg that inform underlying mechanisms linking DNA methylation differences to both exposure and outcome; the lack of mechanistic information is a critical gap in our understanding of most EWAS candidate biomarkers that limits their translational utility^5^.

**Table 1.** Descriptive statistics of age- and sex-matched study participants with either high (>10 μg/g) or low (<1 μg/g) total hair mercury (THg). THg is a proxy for methylmercury (MeHg) exposure. Characteristics of this subset (microarray study population) are compared to the larger biomarker study population. The “Native” column reports on the number of group participants living in an indigenous community (as outlined in Weinhouse *et al* 2021^6^).

|  |  |  |  |
| --- | --- | --- | --- |
| Biomarker study population | N |  | 274 |
|  | Age (median (range)) |  | 32 (18-55) |
|  | Sex |  |  |
|  |  | Male | 91 |
|  |  | Female | 183 |
|  | Native (%) |  | 33% (89/274) |
|  | Total hair mercury (µg/g) (median (range)) |  | 2.2 (0.18-22.7) |
| Microarray study population | High MeHg |  |  |
|  | N |  | 16 |
|  | Age (median (range)) |  | 32 (24-40) |
|  | Sex |  |  |
|  |  | Male | 8 |
|  |  | Female | 8 |
|  | Native (%) |  | 69% (11/16) |
|  | Total hair mercury (µg/g) (median (range)) |  | 12.4 (10-22.7) |
|  | Low MeHg |  |  |
|  | N |  | 16 |
|  | Age (median (range)) |  | 30 (24-43) |
|  | Sex |  |  |
|  |  | Male | 7 |
|  |  | Female | 9 |
|  | Native (%) |  | 0.06% (1/16) |
|  | Total hair mercury (µg/g) (median (range)) |  | 0.48 (0.29-1.06) |

**Table 2.**

| Table 2. |  |  |  |  |  |
| --- | --- | --- | --- | --- | --- |
| CG ID | Mean High MeHg | Mean Low MeHg | Mean Difference | FDR | Gene annotation |
| cg02454636 | 0.70 | 0.74 | -0.04 | 3.98E-05 | SLC7A5 |
| cg02389264 | 0.75 | 0.79 | -0.05 | 3.98E-05 | NA |
| cg22007216 | 0.83 | 0.88 | -0.05 | 4.53E-05 | DCHS2 |
| cg11418303 | 0.88 | 0.92 | -0.04 | 2.92E-04 | PTGER3 |
| cg17220548 | 0.77 | 0.81 | -0.04 | 5.52E-04 | NA |
| cg11802689 | 0.10 | 0.03 | 0.07 | 2.96E-03 | GALNT14 |
| cg01135780 | 0.05 | 0.04 | 0.004 | 8.56E-03 | EBF1; LINC02202 |
| cg25777422 | 0.33 | 0.35 | -0.02 | 8.56E-03 | PIP5K1B |
| cg18944274 | 0.88 | 0.86 | 0.02 | 3.29E-02 | NA |

Based on our results, we propose that future EWAS studies prioritize exposure variance for candidate biomarker detection, to be followed by replication and biological validation. This goal can be accomplished by focusing EWAS studies in geographic locations with known variance in a given exposure. For example, an EWAS for arsenic exposure is best performed in a population of Bangladeshi individuals with high and variable exposure to arsenic due to ingestion of contaminated groundwater^7^. This approach would also improve the ethnic and genetic diversity of EWAS cohorts; most EWAS are currently performed in study populations from Europe or of European descent^5,8^. The DNA methylation biomarker candidates derived from these smaller, targeted studies could be used to generate custom sequencing panels or microarrays specific to each environmental pollutant, dietary component or stressor of interest for use in populations with more limited exposure variance. Custom panels or arrays would incorporate many fewer CpG sites than current epigenome-wide platforms, which would reduce the number of statistical comparisons during the analysis phase^5^. Our proposed approach focused on targeted EWAS that maximize exposure variance would move the field forward by improving epigenetic biomarker detection with efficient use of research resources.

## III. Methods

### Sample population

This study leverages a larger mercury exposure assessment study in communities around the Amarakaeri Communal Reserve in Madre de Dios, Peru^6^. This reserve is bordered on the east by heavy artisanal and small-scale gold mining activity (ASGM), a form of mining that uses large inputs of elemental mercury and contaminates local fish with methylmercury^6^. Residents of these communities are exposed primarily to methylmercury by consuming methylmercury-contaminated fish, although other dietary sources of MeHg and environmental exposure to inorganic mercury via mercury-gold amalgam burning are possible additional sources^6^. We previously quantified total mercury levels in proximal 2-centimeter segments of head hair, which represents ∼2-3 months’ growth^6,9^. For populations in this region, methylmercury is the dominant form of mercury in scalp hair^9^. Thus, total mercury level in this hair segment length approximates primarily methylmercury exposure over the prior 2-3 months^6,9^. For this study, we selected individuals from a subset of the parent study population for which we collected biomarker data from biological samples, including total hair mercury and DNA extracted from PAXgene blood tubes (**Table 1**); this biomarker sub-study is representative of the larger parent study^6^. From this biomarker study subset, we selected 16 adults with high chronic methylmercury exposure (defined as >10 μg/g total hair mercury) and 16 adults with low chronic exposure (defined as <1 μg/g total hair mercury), matched on age and sex^6^ (**Table 1**).

### DNA extraction

For both DNA methylation and mtDNA analyses, 8.5 mL of whole blood was collected in PAXgene Blood DNA Tubes (Qiagen, 761115) which contain 2 mL of a proprietary additive that prevents coagulation of the blood and preserves genomic DNA. Tubes were stored for no more than four hours at room temperature, transferred to a −20°C freezer for a period of four to seven days, and finally transferred to −80° until being shipped on dry ice to Duke University where they were stored at −80°C until DNA was isolated. For DNA isolation, the frozen whole blood samples were thawed in a 37°C water bath for 15 minutes and then immediately processed. PAXgene Blood DNA kits (QIAGEN, 761133) were used according to the manufacturer’s instructions to extract high molecular weight DNA from tissue (not cells).

### DNA methylation

We assessed genome-wide DNA methylation using Illumina Infinium MethylationEPIC BeadChips. We analyzed DNA methylation microarray data in R using the standard pipeline in RnBeads 2.0^10^. Briefly, this pipeline includes quality control via analysis of array control probes and genotyping probes; pre-normalization filtering of probes containing SNPs, or containing high detection p-values (using the Greedycut algorithm); normalization using the dasen method^11^, which includes background correction, between-array normalization applied to Type I and Type II probes separately and no dye-bias correction; post-normalization filtering of probes located on sex chromosomes; and imputation of missing data using the mean methylation level for a given CpG site across all samples^10,12^. We estimated cell type proportions within whole blood samples using the *estimateCellCounts* function in the *minfi* package^13^ and estimated pairwise associations between age, sex, binary MeHg exposure, and cell type proportion variables. In addition, we estimated epigenetic age using the MethylAger algorithm, which is incorporated into the RnBeads pipeline, and compared to chronological age, to account for different epigenetic age pacing between younger and older age groups^14^. Using the MethylAger tool, we used a pre-defined age predictor developed from training methylation datasets from multiple, publicly available studies as a reference; these training datasets include Infinium 27K BeadChip (N=2,286 from 6 studies), Infinium 450K BeadChip (N=1,866 samples from 20 studies from Gene Expression Omnibus or The Cancer Genome Atlas), and Reduced Representation Bisulfite Sequencing data (N=232 samples of German origin). Training datasets include majority European samples (datasets are listed at https://github.com/epigen/RnBeads_web/blob/master/ageprediction.html.)

We conducted differential methylation analysis on the site and region level between high and low MeHg groups (based on a binary MeHg variable) adjusted for sex, age, community and estimated proportions of the following cell types: CD8+ T cells, CD4+ T cells, B cells, natural killer cells, monocytes, granulocytes. RnBeads computed p-values and adjusted p-values (using the Benjamini-Hochberg false discovery rate (FDR) correction for multiple comparisons) on the site and region levels using hierarchical linear models from the *limma* package and fitted using an empirical Bayes approach on derived M-values. Then, RnBeads assigned ranks to differentially methylated sites based on three criteria: 1) the difference in mean methylation, 2) the quotient in mean methylation, and 3) a statistical test (results from the multivariable regression). A combined rank was computed based on the maximum rank among these three metrics (the lower the rank, the greater the evidence for differential methylation). Differentially variable sites were computed using the *diffVar* method from the *missMethyl* R package^15^: 1) the mean differences in means across all sites in a region between high and low MeHg groups, 2) the mean of quotients in mean methylation, and 3) the combined p-value from all site p-values in the region. Each region was assigned a combined rank based on the maximum rank among these three metrics. Regions are defined as belonging to one of four genetic context categories: genomic tiling (5000 bp genomic windows with dense probe coverage), CpG islands (Ensembl annotations), gene promoters (1.5 kb upstream and 0.5 kb downstream of transcription start sites), and genes (whole loci from transcription start sites to transcription end sites)^10^. Differential variability on the region level was computed similarly to differential methylation on the region level, using the mean of variances, log-ratio of the quotient of variances, and p-values from the differentiality test to compute ranks. We conducted a Gene Ontology (GO) enrichment analysis of the top 1000 ranked sites and regions using a hypergeometric test^16^, as well as a Locus Overlap Analysis (LOLA) enrichment^17^ using Fisher’s exact tests to derive ranked enrichments in functional genomic and epigenomic annotations from the following reference databases: cistrome_cistrome, cistrome_epigenome, codex, encode_segmentation, encode_tfbs, Sheffield_dnase, and uscs_features.

### Mitochondrial DNA damage and copy number

We assessed mitochondrial DNA copy number (mtDNA CN) and mitochondrial DNA damage (mtDNA damage) as follows. DNA was quantified using PicoGreen (ThermoFisher P7589) with a standard curve of a HindIII digest of lambda DNA (Invitrogen 15612-013) as described^18^. Samples were then diluted to 3 ng/μL in 0.1X TE buffer for use in long amplicon Polymerase Chain Reaction (LA-PCR) and real time PCR assays. We measured mtDNA damage using an established long-range qPCR assay that evaluates whether DNA lesions are present that can halt or slow DNA polymerase progression during PCR amplification. This assay’s primers amplify an 8.9 kb fragment from mtDNA. Samples with greater loads of DNA damage yield fewer PCR products. For each mtDNA damage qPCR reaction, we used 15 ng DNA template, 0.4 µM each of forward (5’-TCT AAG CCT CCT TAT TCG AGC CGA-3’) and reverse (5’-TTT CAT CAT GCG GAG ATG TTG GAT GG-3’) primers, nuclease-free water, and LongAmp Hot Start Taq 2× Master Mix (New England Biolabs), as described^18^. We amplified this product under the following conditions: an initial denaturation step of 2 min at 94°C, 21 cycles of denaturation at 94°C for 15 seconds and annealing at 64°C for 12 minutes, with a single final extension step at 72°C for 10 minutes. We quantified qPCR products using Picogreen dye in a 96-well plate reader as described^18^. We calculated DNA lesion frequency for mtDNA following a Poisson equation [f(x) = e^-lλ^ λ^x^/x!], where λ is the average lesion frequency in the reference template (i.e., the zero class; x=0, f(0) = e^-λ^), as previously described^19^. We compared amplification of mtDNA in people with high hair mercury (A_HIGH_) to amplification of mtDNA in people with low hair mercury (A_LOW_) with a relative amplification ratio (A_HIGH_/A_LOW_). We defined the DNA lesion frequency as λ = - ln(A_HIGH_/A_LOW_). We calculated lesion frequency per base pairs (bp) of mtDNA by adjusting for amplicon size and normalizing amplification of the long mtDNA fragment to the short mtDNA fragment that reflects mtDNA CN per cell^18^.

We measured mtDNA CN using an established short-range, real-time, standard curve-based qPCR assay that is specific to mtDNA. We prepared serial dilutions of a plasmid containing a 107-base fragment of the mitochondrial tRNA-Leu(UUR) gene to create a standard curve to then calculate absolute mtDNA CN, as previously described^18^. We evaluated associations between MeHg and mtDNA CN or mtDNA damage with tests of correlation (**Fig. 2B, Supplemental Fig. 2**), as well as multivariate regression models, adjusted for age, sex, and cell type proportions.

## IV. Results

### Differential DNA methylation at the site and region levels

We observed nine CpG sites with differential DNA methylation between high and low MeHg groups that remained significant after adjusting for multiple comparisons (FDR <0.05) (**Table 2**). Most of these significant CpG sites are near or within genes that are functionally linked to methylmercury toxicity. The top hit CpG site (cg02454636) is located within an intron of the *SLC7A5* gene, which encodes the L-type amino acid transporter 1 (LAT1). LAT1 is one of two transporters that are responsible for MeHg’s ability to distribute throughout the body and exert toxicity^20–22^. MeHg forms a complex with L-cysteine (derived from glutathione); this complex is a structural mimic for large amino acids like methionine and can be transported into protein-rich tissue, including brain and muscle, via LAT1 and LAT2^20–22^.

Three additional hits suggest that the PI3K/Akt/mTOR signaling pathway is differentially regulated in individuals with high MeHg exposure. Two of the additional significant CpG sites, cg01135780, located within the long non-coding RNA *LINC02202*, and cg25777422, located within a promoter element in the first exon of the *PIP5K1B* gene, are implicated in PI3K signaling. PI3K is an upstream regulator of mTOR-triggered autophagy and is involved in MeHg-induced oxidative stress in neurons^23–25^ and MeHg-inhibited insulin secretion in pancreatic β cells^26^. Relatedly, cg11802689 is located within an intronic enhancer of the *GALNT14* gene; GALNT14 is a glycosyltransferase that regulates apoptosis through mTOR^27^.

Three more differentially methylated CpG sites (cg11418303, cg22007216, cg01135780) are associated with known pathways of MeHg toxicity or known health outcomes related to MeHg exposure. Cg11418303 is located within an intronic enhancer in the *PTGER3* (*prostaglandin E receptor 3*) gene. This receptor is one of four that interacts with prostaglandin E2; this receptor is elevated in islets of diabetic db/db mice and its pharmacologic blockage enhances pancreatic β cell proliferation and improves Nrf2-mediated protection against MeHg-induced oxidative stress. Cg22007216 is located within an intron in the *DCHS2* (*dachsous cadherin related 2*) gene, which is predicted to enable calcium ion binding; MeHg exerts toxicity partially through dysregulation of calcium ion signaling^28,29^. In addition to its annotation with LINK02202, cg01135780 is annotated with the gene *EBF1* (*early B-cell factor 1*), which regulates expression of proteins important for B cell differentiation and function; MeHg is immunotoxic and has been reported to suppress B cell function and antibody formation^30,31^.

The remaining three differentially methylated CpG sites were not annotated in the Illumina manifest (**Table 2**) and are located near several genes each, leaving their gene regulatory potential unclear. We did not observe any differentially methylated regions after adjustment for multiple comparisons. Tables with complete data are available in the Gene Expression Omnibus (GSE207443).

### GO enrichment in differential and differentially variable DNA methylation

In addition to testing for differentially methylated sites and regions, we performed Gene Ontology (GO) enrichment analysis to further explore the biological signal in our dataset. GO enrichment analysis leverages the GO Consortium’s curated, logical hierarchy of gene sets and their functional annotations to identify genes enriched within discovery datasets^16^. These gene sets are curated into groupings, or GO “terms”, within three categories: Biological Processes, Molecular Functions, and Cellular Components^16^. We observed enrichments of GO terms for regions in genes and promoters only, and no enrichments for genomic tiling regions or CpG islands (region types are defined in Methods). We observed 112 Biological Process (BP) GO terms enriched in gene regions and 51 terms enriched in promoter regions with hypomethylated DNA in high vs. low MeHg groups (using the 1000 best ranking regions, as described in Methods, all with p≤0.01) (**Supplemental Tables S1-2**, selected terms related to the most commonly enriched biological themes in **Table 3**). We observed 46 BP GO terms enriched in gene regions and 51 terms enriched in promoter regions with hypermethylated DNA in high vs. low MeHg groups (using a cutoff of combined rank among the 1000 best ranking regions, all with p≤0.01) (**Supplemental Tables S3-4**, selected terms in **Table 4**). In addition, we saw 70 BP GO terms in genes and 128 BP GO terms in promoters with hypervariable DNA methylation between exposure groups, using the same cutoffs (**Supplemental Tables S5-6**, selected terms in **Table 5**), as well as 33 terms in gene regions and 43 terms in promoter regions with hypovariable DNA methylation between exposure groups (**Supplemental Tables S7-8**). Most enriched GO terms were related to immune response, with a particular focus on the innate immune response/inflammation (**Tables 3-5, Supplemental Tables S1-S8**).

**Table 3.** Selected Biological Process terms from the top 1000 ranked terms in the Gene Ontology (GO) database enriched in genes and promoters hypomethylated in Peruvians with high vs. low MeHg exposure. The full list of enriched GO terms can be found in Supplemental Tables S1 and S2.

|  |  |  |
| --- | --- | --- |
| Genes |  |  |
|  | GO:0002758 | Innate immune response-activating signal transduction |
|  | GO:0045089 | Positive regulation of innate immune response |
|  | GO:0006954 | Inflammatory response |
|  | GO:1903555 | Regulation of tumor necrosis factor superfamily cytokine production |
| | GO:1901224 | Positive regulation of NIK/NFκB signaling |
| | GO:0032651 | Regulation of interleukin-1β production |
|  | GO:0032733 | Positive regulation of interleukin-10 production |
|  | GO:0032612 | Interleukin-1 production |
| | GO:0050725 | Positive regulation of interleukin-1β biosynthetic process |
|  | GO:0032755 | Positive regulation of interleukin-6 production |
| | GO:0032729 | Positive regulation of interferon $\gamma$ production |
|  | GO:0042119 | Neutrophil activation |
|  | GO:0002275 | Myeloid cell activation involved in immune response |
|  | GO:0002444 | Myeloid leukocyte-mediated immunity |
|  | GO:0034241 | Positive regulation of macrophage fusion |
|  | GO:0002351 | Serotonin production involved in inflammatory response |
|  | GO:0060585 | Positive regulation of prostaglandin-endoperoxide synthase activity |
|  | GO:0070101 | Positive regulation of chemokine-mediated signaling pathway |
|  | GO:0002371 | Dendritic cell cytokine production |
| | GO:2000516 | Positive regulation of CD4+, $\alpha\beta$ T cell activation |
| | GO:0046641 | Positive regulation of $\alpha\beta$ T cell proliferation |
| | GO:0046637 | Regulation of $\alpha\beta$ T cell differentiation |
|  | GO:0042088 | T-helper 1 type immune response |
|  | GO:0038156 | Interleukin-3 mediated signaling pathway |
| Promoters |  |  |
|  | GO:0050830 | Defense response to Gram-positive bacterium |
|  | GO:0050829 | Defense response to Gram-negative bacterium |
|  | GO:0043303 | Mast cell degranulation |
|  | GO:0032762 | Mast cell cytokine production |
|  | GO:0002548 | Monocyte chemotaxis |
|  | GO:0002315 | Marginal zone B cell differentiation |

**Table 4.** Selected Biological Process terms from the top 1000 ranked terms in the Gene Ontology database enriched in genes and promoters hypermethylated in Peruvians with high vs. low MeHg exposure. The full list of enriched GO terms can be found in Supplemental Tables S3 and S4.

|  |  |  |
| --- | --- | --- |
| Genes |  |  |
|  | GO:0002227 | Innate immune response in mucosa |
|  | GO:0050830 | Defense response to Gram-positive bacterium |
|  | GO:0061844 | Antimicrobial humoral immune response mediated by antimicrobial peptide |
|  | GO:0002548 | Monocyte chemotaxis |
| | GO:2001201 | Regulation of TGF-β secretion |
|  | GO:0038111 | Interleukin-7-mediated signaling pathway |
|  | GO:0098760 | Response to interleukin-7 |
| Promoters |  |  |
|  | GO:1900246 | Positive regulation of RIG-1 signaling pathway |
|  | GO:0038111 | Interleukin-7-mediated signaling pathway |
|  | GO:0098760 | Response to interleukin-7 |

**Table 5.**
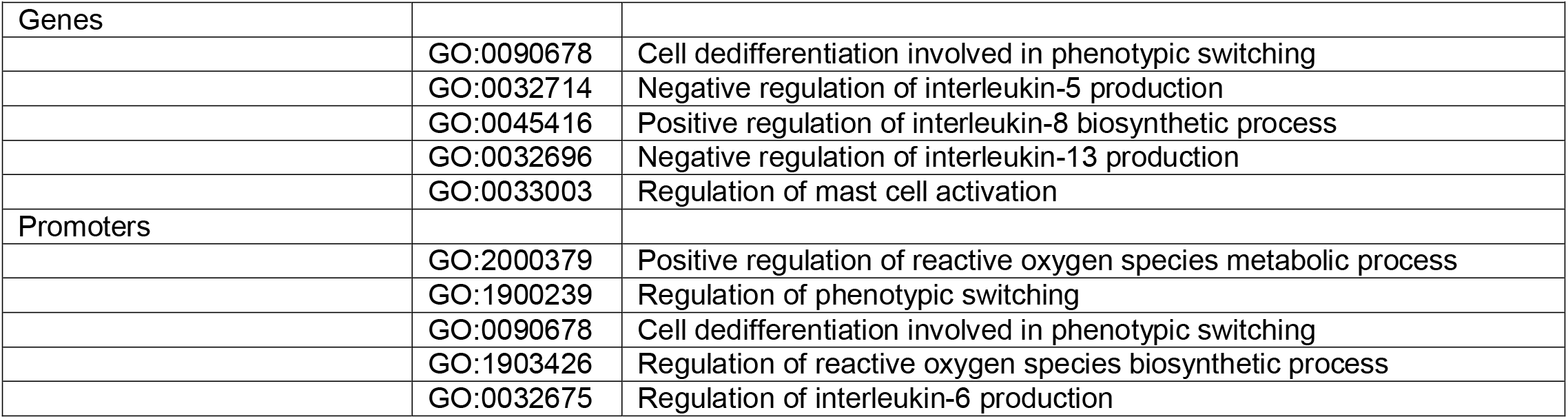
Selected Biological Process terms from the top 1000 ranked terms in the Gene Ontology database enriched in genes and promoters with hypervariable DNA methylation in Peruvians with high vs. low MeHg exposure. The full list of enriched GO terms can be found in Supplemental Tables S5 and S6.

### LOLA enrichment in differential and differentially variable DNA methylation

To complement the gene-centric GO enrichment analysis, we additionally performed Locus Overlap Analysis (LOLA) to identify regulatory regions within our dataset enriched for functional genomic and epigenomic annotations^17^. Using the top 1000 ranked regions, we observed enrichments of LOLA annotations in genomic tiling regions and CpG islands only, and no enrichments for genes or promoters. Most enriched annotations were for binding sites for transcription factors involved in hematopoiesis and immune response in regions hypomethylated in high vs. low MeHg groups (**Fig. 1**, **Supplemental Figs. 3-13**). The second most common signal in our LOLA results was for general repression in regions hypomethylated in high vs. low MeHg groups (**Supplemental Figs. 3-13**). In particular, we observed enrichment in repressive signals in hypomethylated regions in high vs. low MeHg groups, which suggests reactivation of repressed regulatory regions (**Supplemental Figs. 3-13**). Signals of gene repression include repressive histone modifications (e.g., H3K27me3) and loss of methylation (indicating binding and activation) in regions associated with binding of proteins that deposit repressive histone modifications (e.g., polycomb repressive complex components EZH2^32^ and SUZ12^33^) or remove activating histone modifications (e.g., SMARCA4, a component of the SWI/SNF chromatin remodeling complex^34^, which recruits histone deacetylase repressor complexes^35^).

**Figure 1.**
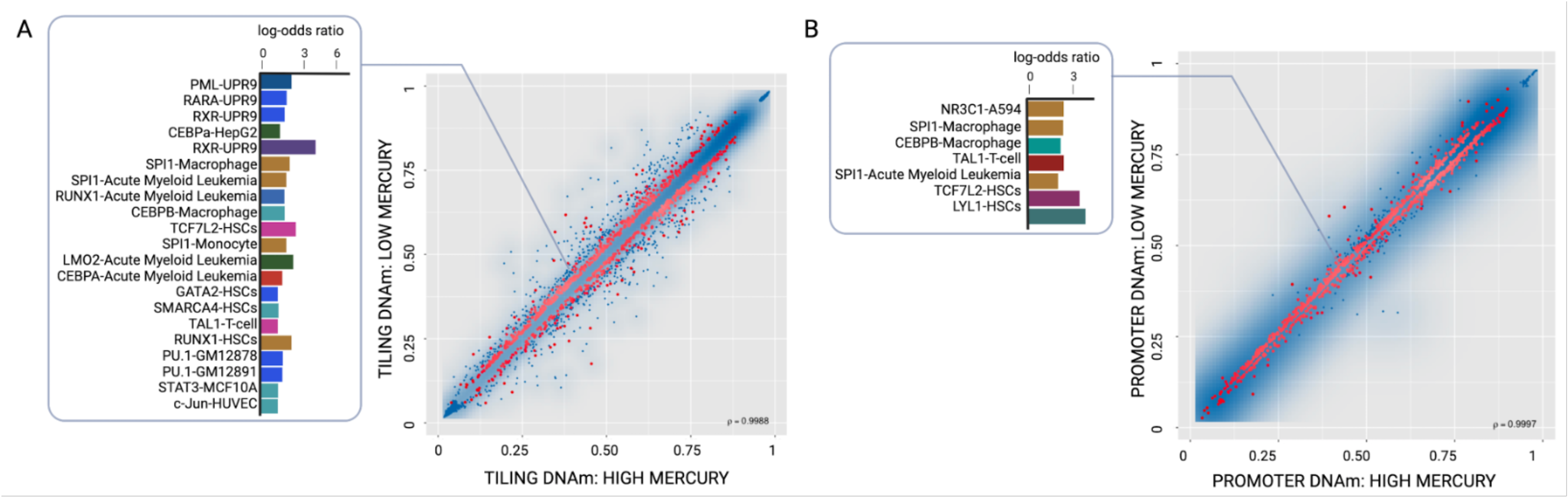
Transcription factor binding site enrichments in genomic regions hypomethylated in individuals with high methylmercury exposure. Scatterplot for differentially methylated (A) genomic tiling regions and (B) promoter regions. Color transparency corresponds to point density; the 1% of points in the sparsest population plot regions are drawn explicitly. Red colored points represent the 1000 best ranking regions; the linked barplots represent enrichments within these top ranked data points. Barplots showing selected log-odds ratios (p<0.01) from LOLA enrichment analysis for (A) genomic tiling and (B) promoter regions that are hypomethylated in Peruvian study participants with high (>10 μg/g) vs. low (<1 μg/g) total hair mercury, a proxy for methylmercury exposure.

### Predicted epigenetic age, cell type proportions, and mitochondrial endpoints

In addition to testing for differentially methylated CpG sites, we evaluated the utility of a small EWAS in a population with high exposure variance to inform other commonly used biomarkers. Specifically, we computed predicted epigenetic age, sometimes referred to as an “epigenetic clock” biomarker using the machine learning-based MethylAger algorithm, based on age-related changes in DNA methylation in specific CpG sites^36–42^. Accelerated aging, as evident from a discrepancy between chronological age and computed epigenetic age, has been associated with environmental pollutants and disease risk in past studies^36–42^. In our data, the predicted epigenetic ages computed from DNA methylation data were consistently lower than reported chronological age (on average 11 years lower, ranging from 8 to 17 years lower) (**Fig 2A**). Since predicted epigenetic age was highly correlated with reported chronological age (R^2^=0.86) (**Fig. 2A**), these results indicate systematically lower age by the MethylAger algorithm in our dataset. In addition, we did not observe any association between predicted epigenetic age and MeHg exposure (**Fig. 2B**). In pairwise tests of association between proportions of different immune cell types with mercury exposure, only monocyte proportion was associated with binary MeHg with higher proportion of monocytes in the high mercury group (Wilcoxon test p=8.7×10^-5^) (**Fig. 2B, Supplemental Fig. 1**). Neither continuous nor binary total hair mercury was associated with mtDNA damage (R^2^=7E-05) (**Fig. 2C**) or mtDNA CN (R^2^=0.0028) (**Fig. 2D**). In addition, mtDNA damage was not highly correlated with mtDNA CN (R^2^=0.07) (**Supplemental Fig. 1**). We observed a broad distribution of both mtDNA damage and mtDNA CN biomarkers in both high and low MeHg exposure groups (**Fig. 2C-D**).

**Figure 2.**
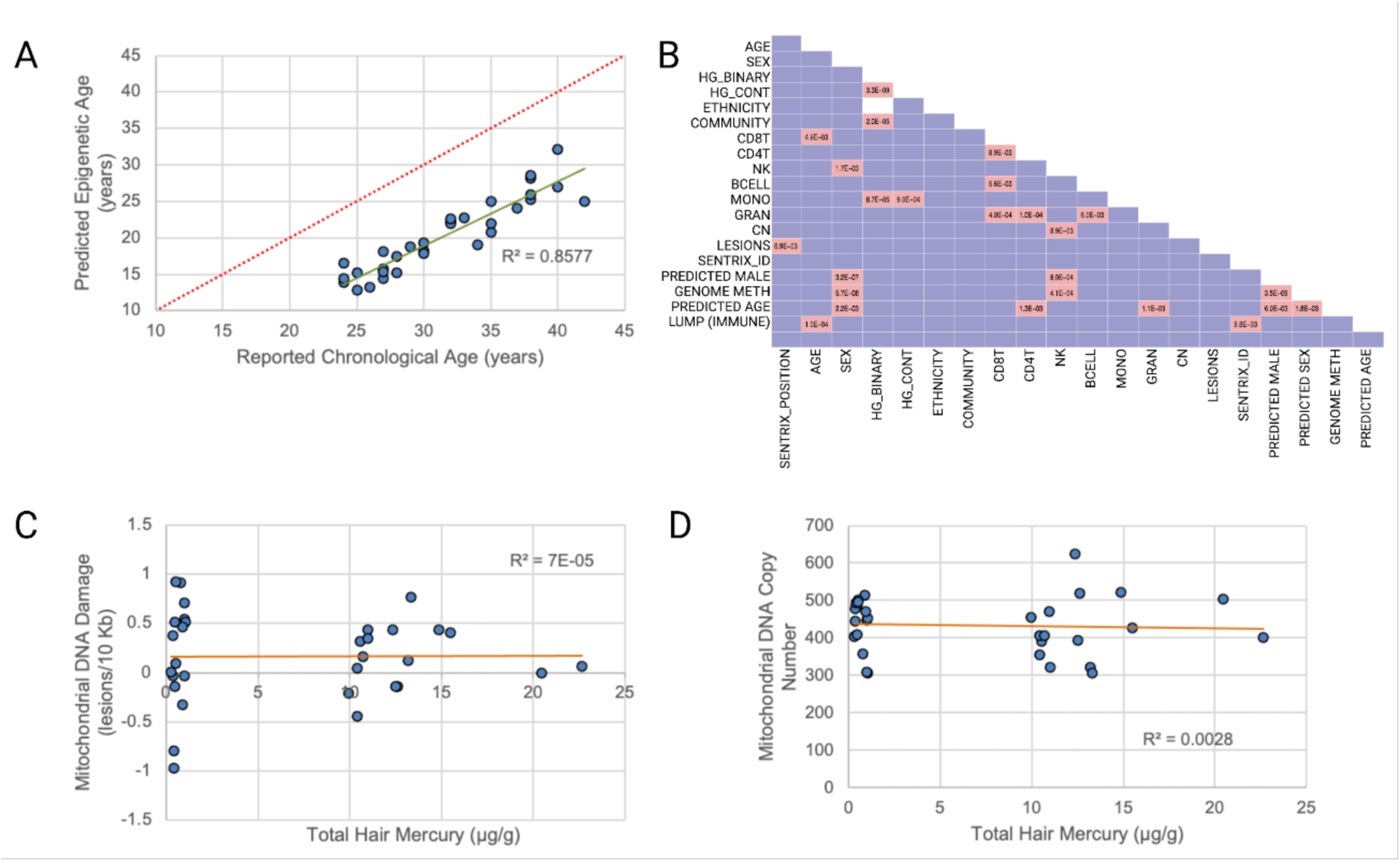
Tests of association for epigenetic age, cell type proportions and mitochondrial DNA biomarkers with methylmercury exposure in Peruvian individuals. (A) Association between predicted epigenetic age and reported chronological age. (B) Pair-wise associations between covariates, including white blood cell type proportions (as estimated by DNA methylation profiles), and methylmercury exposure (estimated by total hair mercury). (C) Association between mitochondrial DNA damage and total hair mercury levels. (D) Association between mitochondrial DNA copy number and total hair mercury levels.

## V. Discussion

In this study, we provide proof-of-principle that EWAS with large exposure variance can yield candidate DNA methylation biomarkers and substantial data informing underlying biological mechanisms, even in very small sample sizes. Specifically, we identified nine differentially methylated CpG sites, including a CpG site in an intronic enhancer of the critical transporter LAT1, in Peruvian individuals with high (>10 μg/g) vs. low (<1 μg/g) total hair mercury (a proxy for methylmercury exposure), matched on age and sex. In addition, our GO term and transcription factor motif enrichment analyses provided highly informative signals of MeHg exposure, including signaling pathways linked to immune system response, type 2 diabetes risk, glucocorticoid receptor-mediated neurotoxicity, and PUFA-mediated protection from MeHg toxicity. Notably, we did not find similarly strong signals in other commonly measured biomarkers of environmental chemical exposure, including mitochondrial DNA damage or copy number, and epigenetic age, indicating that these biomarkers are less well understood and should be measured in small samples with caution.

The primary signal in our GO enrichment data was of a clear immune phenotype in response to MeHg exposure. The human immune response comprises general innate responses as well as antigen-specific adaptive responses^43^. Both innate and adaptive immune responses include a humoral component (circulating chemical effectors, including cytokines and chemokines) and a cell-mediated component (including general neutrophil and macrophage^44^ responses and both general^45,46^ and antigen-specific^46^ B-cell and T-cell action). Most pathways that were enriched in hypomethylated regions in genes and promoters in high MeHg-vs. low MeHg-exposed Peruvians reflect innate immune response activation (**Table 2, Supplemental Tables S1-2**). Loss of DNA methylation in promoters and genes are generally associated with gene activation^47^, implying activation of these innate immune pathways in response to MeHg exposure. This innate response includes classic neutrophil and macrophage activation^48,49^, as well as mast cell release of serotonin^50^ and eicosanoids like prostaglandins^51^ (**Tables 2 and 4, Supplemental Tables S1-S2**). In addition, we observed evidence of immune responses in several T- and B-cell subtypes (**Tables 2 and 4, Supplemental Tables S1-S2**). Both T-cells and B-cells develop effector cells in response to specific antigens, including DAMPs^52^. A subset of each effector cell subtype is retained as memory cells following immune response resolution^53^. T-cell responses are generally divided into cytotoxic (CD8^+^ T-cells) and helper (CD4+ T-cells) responses; CD4^+^ responses are further subdivided into T-helper type 1 (Th1) and T-helper type 2 (Th2) responses^54^. Th1 responses promote inflammation and, if uncontrolled, cause autoimmunity and tissue damage due to chronic inflammation^55^. Th2 responses include anti-inflammatory cytokines, as well as eosinophilic (e.g., IgE- and histamine-mediated signaling), that counterbalance Th1 responses^55^. Here, we observed a clear CD4^+^ response, including enrichment of hypomethylated genes among pathways related to both Th1 (inflammatory cytokines and chemokines: interleukin-1 (IL-1), interleukin-1β (IL-1 β), interleukin-6 (IL-6), interleukin-8 (IL-8), tumor necrosis factor α (TNFα), macrophage-activating interferon γ (IFN γ)) (**Tables 2 and 4, Supplemental Tables S1-S2 and S4-S5**) and Th2 (anti-inflammatory interleukin-10 (IL-10), eosinophil activation, interleukin-5 (IL-5), interleukin-13 (IL-13), B-cell isotope switching) (**Supplemental Tables S1-S2 and S4-S5**). Importantly, the Th1 response is most evident in our GO enrichments of differential mean DNA methylation (**Tables 2 and 3, Supplemental Tables S1-S4**) and the Th2 signal is clearest in GO enrichments of hypervariable DNA methylation (**Table 4, Supplemental Tables S4-S5**). These results suggest that MeHg induces a similar Th1 response in most individuals, but that some individuals mount a stronger balancing Th2 response than others. This result suggests a mechanism by which MeHg-exposed individuals who exhibit Th1-dominant signaling with little Th2 counterbalance may be at higher risk of developing autoimmunity. Therefore, DNA methylation at genes and gene promoters associated with Th2 signaling in our dataset are strong candidate epigenetic biomarkers of individual risk of autoimmune response to MeHg.

We observed two additional signals related to the development of autoimmunity. First, our data are consistent with expansion of autoreactive T cells. Autoreactive helper T cell subsets can form in response to self-antigen; T helper type 17 (Th17) cells are most likely to be autoreactive, followed by Th1 cells^56^. Th17 cells are stimulated to differentiate from naïve CD4^+^ T cells by IL-6 and transforming growth factor-β (TGF-β), which stimulate downstream STAT3 signaling^56^. Our data show enrichment in genes involved in all three of these signals (**Tables 2 and 4, Supplemental Tables S1-S4**), indicating an environment conducive to increased Th17 cell production. Regulatory T (Treg) cells provide negative regulation of Th17 cells and suppress autoimmune responses and disease development^57,58^. Treg cells are derived from naïve CD4^+^ T cells when IL-6 and TGF-β levels decrease during resolution of an inflammatory response^56^. In addition, interleukin-7 (IL-7) signaling promotes expansion of the Treg pool^57,58^. The enrichment in hypomethylated genes (which suggests increased activation) involved in IL-6 and TGF-β signaling in response to MeHg, as well as hypermethylation of genes (suggesting decreased activation) involved in IL-7 signaling (**Table 3, Supplemental Tables S3-S4**), are consistent with an expanded pool of autoreactive Th17 cells and a diminished population of Treg cells that suppress autoreactivity in individuals with high MeHg exposure. Second, we observed that hypomethylated regions in high MeHg-exposed were enriched for gene promoters involved in marginal zone B cell differentiation (**Table 2**), which is consistent with activation of the target genes of these promoters. Marginal zone B cells can become autoreactive when co-stimulated by self-antigens and DAMPs^59^, and autoreactive marginal zone B cells can also activate autoreactive T cells^59^. Therefore, in addition to DNA methylation linked to Th2 signaling, hypomethylated enhancers and promoters associated with decreased IL-7 signaling and increased marginal zone B cell differentiation are strong candidates for epigenetic biomarkers that report on inter-individual differences in autoimmune response to MeHg. The hypervariable DNA methylation in multiple inflammatory pathways is strong evidence of population distribution in immune response to MeHg in our dataset, including both high and low responders that carry differential DNA methylation signatures of response (**Supplemental Tables S5-S6**).

The transcription factor signal that we observed in our LOLA enrichments is consistent with the immune phenotype reflected in our GO enrichment data. Specifically, we observed hypomethylation of tiling regions (likely enhancers) and promoters containing binding sites for transcription factors that control differentiation of the macrophages, neutrophils, T-cells and B-cells (**Fig. 1A-B**, **Supplemental Figs. S3-S13).** Broadly, hematopoiesis generates a range of blood cell types, including red (erythrocytes) and white (lymphocytes) cells^60^. Lymphocytes are derived from either myeloid or lymphoid lineages; myeloid precursors differentiate into neutrophils and monocytes/macrophages and lymphoid precursors develop into B-cells and T-cells^60^. Spi1/PU.1 is a master regulator of hematopoiesis that directs differentiation within both myeloid and lymphoid lineages through varying concentration (e.g., low in multipotent precursors, high in mature B-cells and macrophages) and co-activator partners^61^. During early hematopoiesis, Spi1/PU.1 interacts with factors GATA-2, CEBPα/β and c-Jun to drive white blood cell differentiation^61,62^. In the presence of STAT3 (**Fig. 1A**) and interleukin-3 signaling (**Supplemental Table S1**), cells further develop into neutrophils and macrophages^63^. In contrast, RUNX1^64,65^, RUNX3^64,65^, TCF3^66^, and TCF12^66^ (**Fig. 1A**) promote T cell lineage commitment. We observed evidence of signaling through additional hematopoietic transcription factors, including LMO2, which is a scaffold protein that enables formation of protein complexes that include components TAL1, LYL1, GATA-2 that act at varying stages of hematopoiesis, primarily early stages^67–70^. These results are supported by an increase in monocyte cell proportion (on average, 6% in high MeHg vs. 4% monocytes in low MeHg p=8.7×10^-5^) in individuals with high MeHg exposure (**Fig. 2B, Supplemental Fig. S2**). In addition to directing development of specific immune cell types, the transcription factors identified in our dataset have relevant roles in innate immune response that we observed in our GO enrichments. For example, the transcription factor c-Fos is a component of the master factor activator protein 1 (AP1) that activates downstream innate immunity^71^. STAT3 mediates cytokine signaling, partly by upregulating c-Fos^71^. BATF is another member of the AP-1 family that dimerizes with Jun proteins and provides negative feedback to AP1 transcription^72^. Last, some of the proteins with binding sites enriched in high vs. low MeHg-exposed individuals regulate chromatin remodeling and transcription. For example, SMARCA4 is a component of the SWI/SNF chromatin remodeling complex^73^.

Some of our results are specifically relevant to our study population. Our data point to a potential mechanistic link between MeHg exposure and type 2 diabetes (T2D). T2D is characterized by persistently high blood glucose levels due to impaired insulin secretion from pancreatic β cells, insensitivity to insulin in peripheral tissues, and increased glucose production in the liver^74^. Several well-established genetic risk factors for T2D are variants in the *transcription factor 7-like 2 (TCF7L2)* gene^75–77^ that drive expression of functional splice isoforms of this gene^76,78^. The protein product of this gene is the high mobility group box-containing transcription factor TCF7L2 which activates Wnt signaling with tissue-specific outcomes^75,79–81^. In pancreatic β cells, human TCF7L2 variants impair normal insulin production and secretion in response to glucose^82,83^; impaired insulin response could lead to T2D, which is supported by the positive correlation between TCF7L2 variant frequency and population T2D risk^84^. In enteroendocrine cells, TCF7L2 may influence T2D susceptibility through its transcriptional regulation of proglucagon, which is the precursor of the insulinotropic peptide hormone glucagon-like peptide 1 (GLP-1)^75^. Together with insulin, GLP-1 regulates blood glucose homeostasis^78^. Our data show that DNA binding sites for TCF7L2 are enriched in tiling regions (likely enhancers) and in gene promoters that are hypomethylated in Peruvians with high vs. low MeHg exposure (**Fig 1A**, **Supplemental Fig. 3**). Loss of DNA methylation in these regions likely reflects binding of TCF7L2 to regulatory binding sites and activation of downstream signaling. If MeHg triggers aberrant TCF7L2 signaling in pancreatic β cells or enteroendocrine cells, in addition to the leukocyte signal observed in this study, then hypomethylation of TCF7L2 enhancers in blood cells may serve as a surrogate tissue biomarker of early MeHg-related T2D risk. MeHg exposure is toxic to pancreatic β cells^85^. However, MeHg is related to T2D risk in some but not all epidemiological studies (reviewed in ^86^). Most notably, cross-sectional analyses in the population-representative National Health and Nutrition Examination Surveys (NHANES) in the United States^87^ and Taiwan^88^ show positive associations between T2D and MeHg exposure. A large prospective human study confirmed this positive association^89^. In contrast, several cross-sectional and prospective human studies report no association^90–92^, or even an inverse association^93^ (attributed to higher consumption of protective dietary nutrients in high exposed groups^93^), between MeHg and T2D risk. These equivocal results suggest population-specific risk profiles. American Indians in the U.S. have higher diabetes risk than do other ethnic groups, which suggests a higher baseline genetic risk in indigenous Peruvians that may be exacerbated by diet and environment^94^. Individuals carrying TCF7L2 risk alleles that develop impaired glucose tolerance show increased conversion of this pre-diabetic state to full T2D onset, as compared to glucose-intolerant non-carriers^95^. These data suggest that MeHg-induced TCF7L2 signaling may pose a greater disease risk in a population with a higher baseline risk for disease.

Some of our results may be more generalizable to health outcomes of MeHg in Western populations. Importantly, our results inform potential biological mechanisms that have not been resolvable in Western epigenetic datasets, possibly due to lower variance in exposure. For example, DNA binding sites for glucocorticoid receptor (GR), encoded by the *NR3C1* gene, are enriched in gene promoters that are hypomethylated in individuals with high vs. low MeHg exposure (**Fig. 1B, Supplemental Fig. S5**) and the GO term “response to dexamethasone” (GO:0071548), which reflects GR activation by dexamethasone, is enriched in hypomethylated genes (in high vs. low MeHg-exposed individuals) (**Supplemental Table S1**). In leukocytes, GR signaling is important for dampening and resolving inflammatory responses^96^. Importantly, in the hippocampus, MeHg exerts neurotoxicity through GR signaling^97^. MeHg binds GR directly and attenuates GR activation by endogenous ligands, leading to decreased GR signaling that contributes to developmental neurotoxicity of MeHg^97^. In addition, rats exposed during development to a complex environmental contaminant mixture containing MeHg showed a dampened ability to reduce serum corticosterone levels following an experimental acute stress event^98^; because GR is responsible for returning corticosterone levels to homeostatic levels in healthy animals, these data provide functional evidence of GR signaling disruption in MeHg-exposed animals^98^. A prior study provides initial evidence that an epigenetic biomarker of MeHg GR inhibition in blood may reflect signaling in brain; specifically, *in utero* mercury exposure to child participants in the Seychelles Child Development Study predicted leukocyte DNA methylation of the *NR3C1* gene^99^. Future work should focus on whether DNA methylation of *NR3C1* gene in blood reflects similar epigenetic profiles at this gene in hippocampus in rodents exposed to environmentally relevant levels of MeHg. If confirmed, DNA methylation at this gene in blood may serve as an actionable and accessible surrogate epigenetic biomarker of MeHg-induced neurotoxicity and whether this signal is specific to prenatal exposure or can occur in adulthood.

Another important finding from our results suggests an epigenetic biomarker for a protective biological response to fish consumption. DNA binding sites for the transcription factors retinoid X receptor (RXR) and retinoic acid receptor α (RARα) are enriched in tiling regions (likely enhancers) that are hypomethylated in Peruvians with high vs. low MeHg exposure (**Fig 1A, Supplemental Figs. S3 and S5**). PUFA found in large, fatty fish, including docosoehexaenoic acid (DHA), activates RXR signaling^100,101^ that triggers downstream antioxidant signaling which protects against MeHg-induced neurotoxicity^102,103^. RXR can form heterodimers with RARα^102^; RXR-RARα signaling is critical for the hippocampus-dependent learning and memory^104^, as well as DHA-augmented fetal neurodevelopment^102^, that is disrupted by early life MeHg exposure^105^. The enrichment for DNA binding sites for transcription factor PML (**Fig 1A, Supplemental Figs. S3 and S5**) in our data likely reflects RXR-RARα signaling, providing further support for activation of this pathway; this signal likely reflects binding sites within the queried database of a cancer fusion gene of PML and RARα that heterodimerizes with RXR and binds to RXR-RARα DNA binding sites^106^. Since human MeHg exposure in Madre de Dios occurs primarily through fish consumption, individuals with the highest MeHg exposure also have the highest fish consumption^6^. Birth cohort data from the high fish- and seafood-consuming populations in the Republic of Seychelles and the Faroe Islands highlight the importance of considering the health benefits of fish consumption, which may outweigh the harms of MeHg exposure in some exposure settings^105,107^. Future work should explore whether epigenetic biomarkers of RXR-RARα activation by fish consumption reflect RXR-RARα in hippocampus, which is the primary target of MeHg neurotoxicity. Validation of a biomarker that reports on how protective fish consumption is for a particular individual is a critical step in providing individualized health recommendations to individuals, particularly those at high risk for harm, like pregnant women and small children.

In addition to differential DNA methylation, we investigated three additional biomarkers that may inform MeHg response in our study participants: epigenetic age and two mitochondrial biomarkers, mtDNA damage and mtDNA CN. We observed that a commonly used epigenetic age algorithm predicted lower epigenetic ages, relative to chronological ages, for all participants in this study (**Fig. 2A**). This result has two possible explanations. The first is that the individuals in our sample have younger epigenetic ages, relative to their chronological ages. If true, this result suggests that our study participants are healthier than the European populations that were used to derive the clocks, possibly due to diet and physical activity differences. The second possible explanation is that the epigenetic age algorithm systematically underestimated chronological age in our sample. If true, this result suggests that current algorithms, which have been trained and tested on datasets from European individuals, may not be generalizable to non-European populations. For algorithms to be more generalizable tools, they should be trained and tested on more diverse datasets. In addition, we observed no relationship between either mitochondrial biomarker and MeHg exposure (**Fig. 2C-D**), as well as promoter hypermethylation of the RIG-1 signaling pathway (**Supplemental Table S4**) through which DAMPs trigger innate immune responses^108^. Although prior papers have not reported clear associations between mitochondrial biomarkers and MeHg exposure, it was unclear whether this lack of signal was statistical or biological^109,110^. The clear signal in immune signaling pathways in our data, coupled with the lack of association between mitochondrial biomarkers and MeHg, indicates that the biological role of mitochondria in the response to MeHg is more complex than previously thought. For example, mitochondrial DAMPs may serve as a signal of self-damage that triggers endogenous suppression of inflammation to promote healing^111^. The high variance in both mitochondrial markers in both high and low exposure groups (**Fig. 2C-D**), as well as hypervariable promoter DNA methylation in pathways involved in ROS production and response (**Table 4**), strongly implies unmeasured source(s) of variation in our population that require study before these biomarkers can be fully realized in population health settings.

It is worth discussing why the primary transcription factors in our LOLA enrichments function during cellular differentiation, even though our study profiled mature, circulating white blood cells. There are two possible explanations for this finding.

The first is that mature cells may carry persistent DNA methylation signatures of past differentiation programs; this possibility is supported by past evidence of similar DNA methylation memories^112^. The second possibility is that these differentiation programs may be reactivated in mature cells to enable dedifferentiation and phenotypic switching between cell types by changing epigenetic programs^113–115^. This possibility is supported by enrichment in our dataset for the GO term “cell dedifferentiation involving in phenotypic switching” (GO:0090678) (**Supplemental Tables S5-S6**).

This study has several important limitations. First, small sample sizes can lead to false positives^116^. Therefore, the candidate DNA methylation biomarkers identified in this study must be replicated and biologically validated before they are incorporated into a custom panel or array for use in other MeHg-exposed populations. Second, our study participants are from a region where residents commonly live in the same villages or towns in which they are born^6^. Therefore, individuals with high adult exposures may have had high developmental exposures, as well. Our results may reflect acute epigenetic responses to MeHg or they may reflect persistent effects of developmental MeHg exposure or a combination of both effects. This ambiguity limits our results’ generalizability to other MeHg-exposed populations. Third, most study participants in the high MeHg group reside in indigenous communities in the Madre de Dios region (**Table 1**), because the highest MeHg exposures accrue to high fish-consuming residents of these native communities^6^. Therefore, we are unable to separate definitively DNA methylation changes due to genetic differences between indigenous and non-indigenous study participants from environmental effects on DNA methylation due to MeHg exposure. However, the differential DNA methylation signal that we observed here largely reflect known biology in MeHg toxicity, which supports a primarily environmental effect, even in the presence of known genetic variation. Fourth, because this study is cross-sectional, there are several limits to results interpretability. For example, our MeHg exposure biomarker reflects only 2-3 months’ prior exposure, which may reflect transient exposure or, alternatively, relatively constant chronic exposure. Therefore, we are unable to assess whether these epigenetic changes reflect responses to short or long exposure durations. In addition, we are unable to assess the persistence, if any, of our observed epigenetic markers. These questions should be assessed in future targeted EWAS cohorts with time-resolved exposures and epigenetic outcomes, to fully realize the potential of small EWAS in highly exposed populations to inform generation of custom panels and arrays for EWAS in populations with lower exposure variance.

## VI. Conclusion

Here, we provide proof-of-principle that EWAS in small samples with high exposure variance may detect candidate DNA methylation biomarkers. Similar EWAS in global populations with known high exposure variance can be leveraged to develop targeted, custom sequencing panels and microarrays for use in EWAS in populations with lower exposure variance.

## Acknowledgments

This work is supported by Hunt Oil Peru LLC (HOEP-QEHSS-140003, W.K.P.), Duke Population Research Institute P2C pilot funds (2P2CHD065563-06 SUB#60P2034949, W.K.P.), NIH grant K01-ES32044-01 (C.W.), a Duke Global Health Institute Postdoctoral Fellowship (C.W. and W.K.P.) and the Duke University Superfund Research Center (P42 ES010356, J.N.M, W.K.P., S.K.M.). The authors acknowledge Ernesto Ortiz and Axel Berky for their roles in data collection for the parent study, as well as study participants and local field workers.

## Data sharing

We have deposited raw data files and sample phenotype data, as well as tables containing differential DNA methylation and differentially variable DNA methylation on both site and region levels, in the Gene Expression Omnibus (GSE207443).

