## Supplemental Figures for "High exposure variance enables candidate biomarker detection in a small EWAS of methylmercury-exposed Peruvian adults"

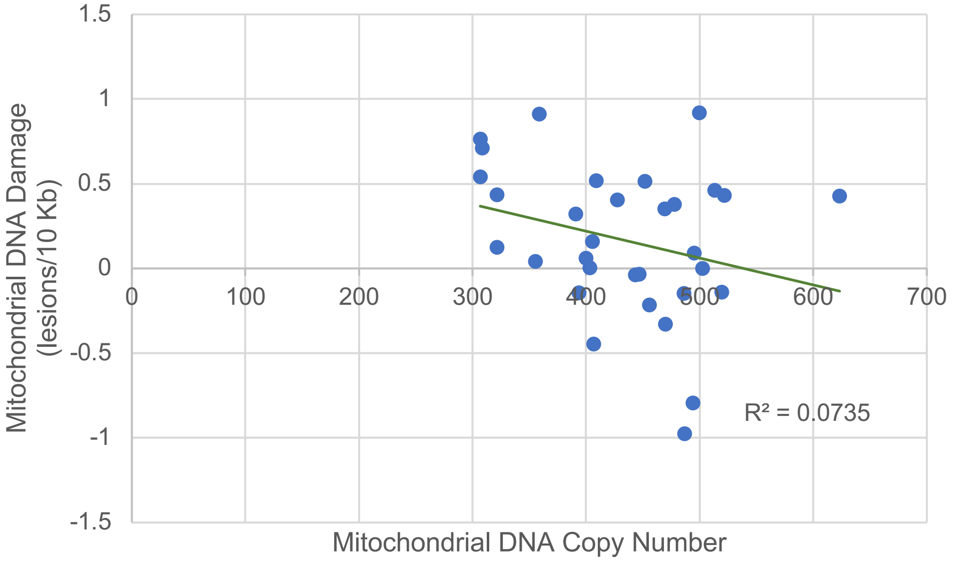


**Supplemental Figure S1. Association between mitochondrial DNA copy number and mitochondrial DNA damage in methylmercury-exposed Peruvian individuals.** Association between mitochondrial DNA copy number and mitochondrial DNA copy number in Peruvian study participants with high (>10 μg/g) vs. low (<1 μg/g) total hair mercury, a proxy for methylmercury exposure.

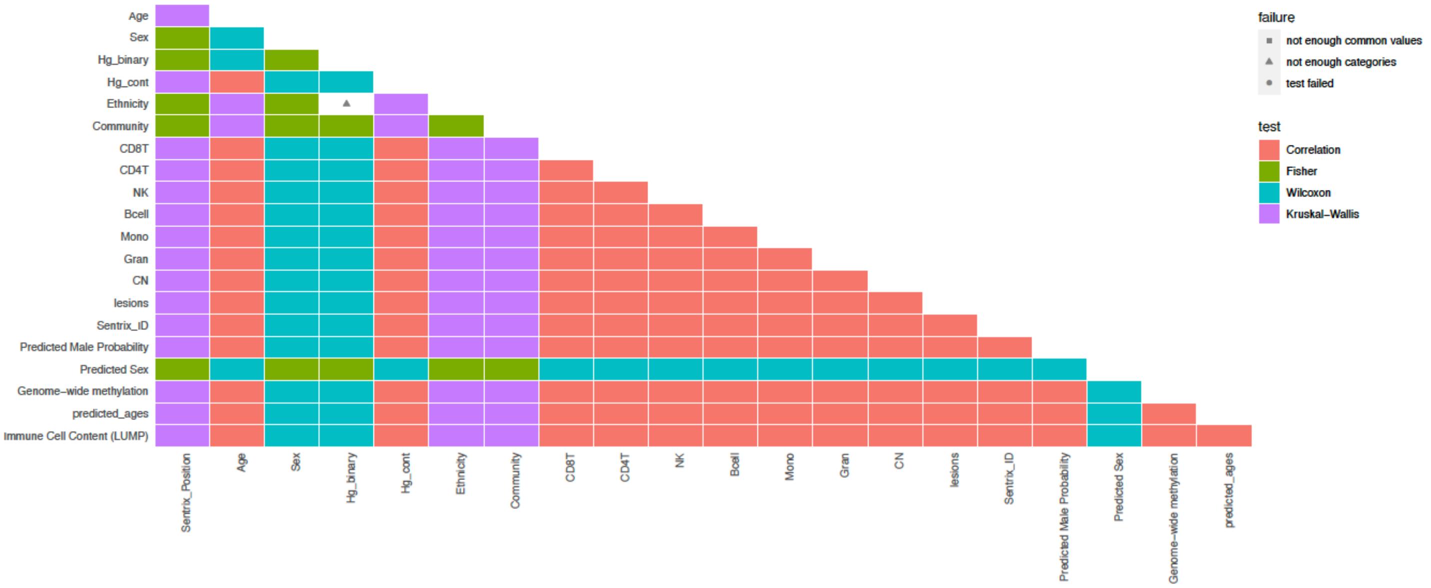


**Supplemental Figure S2. Tests of association for covariates, including cell type proportions, with methylmercury exposure in Peruvian individuals.** Pair-wise associations between covariates, including white blood cell type proportions (as estimated by DNA methylation profiles), and methylmercury exposure (estimated by total hair mercury).

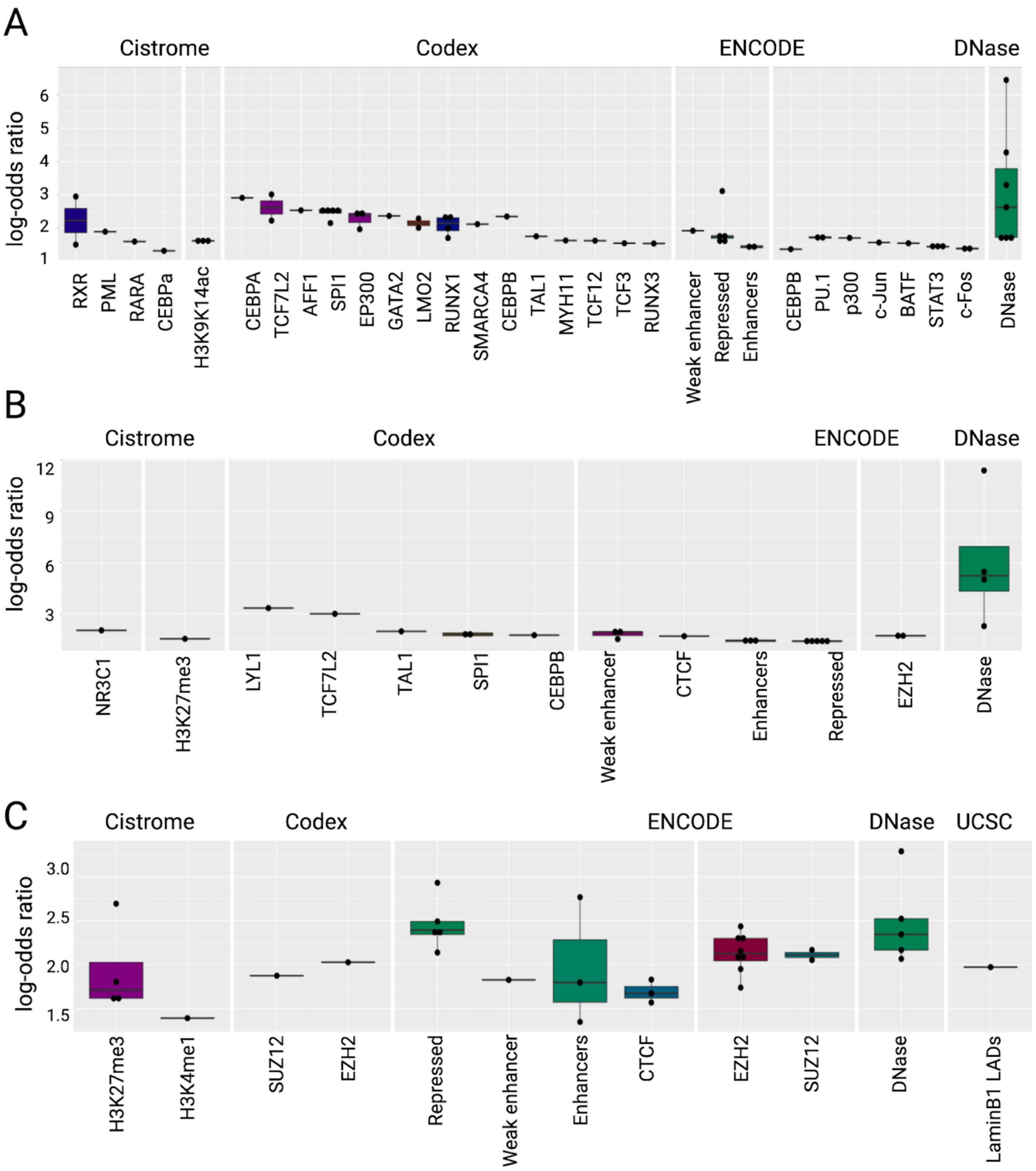


**Supplemental Figure S3. LOLA enrichment in hypomethylated regions in Peruvians with high vs. low methylmercury exposure.** Boxplots showing log-odds ratios ($p<0.01$) for the 1000 best ranking regions in LOLA enrichment analysis for (A) genomic tiling regions, (B) promoters, and (C) CpG islands that are hypomethylated in Peruvian study participants with high (>10 μg/g) vs. low (<1 μg/g) total hair mercury, a proxy for methylmercury exposure.

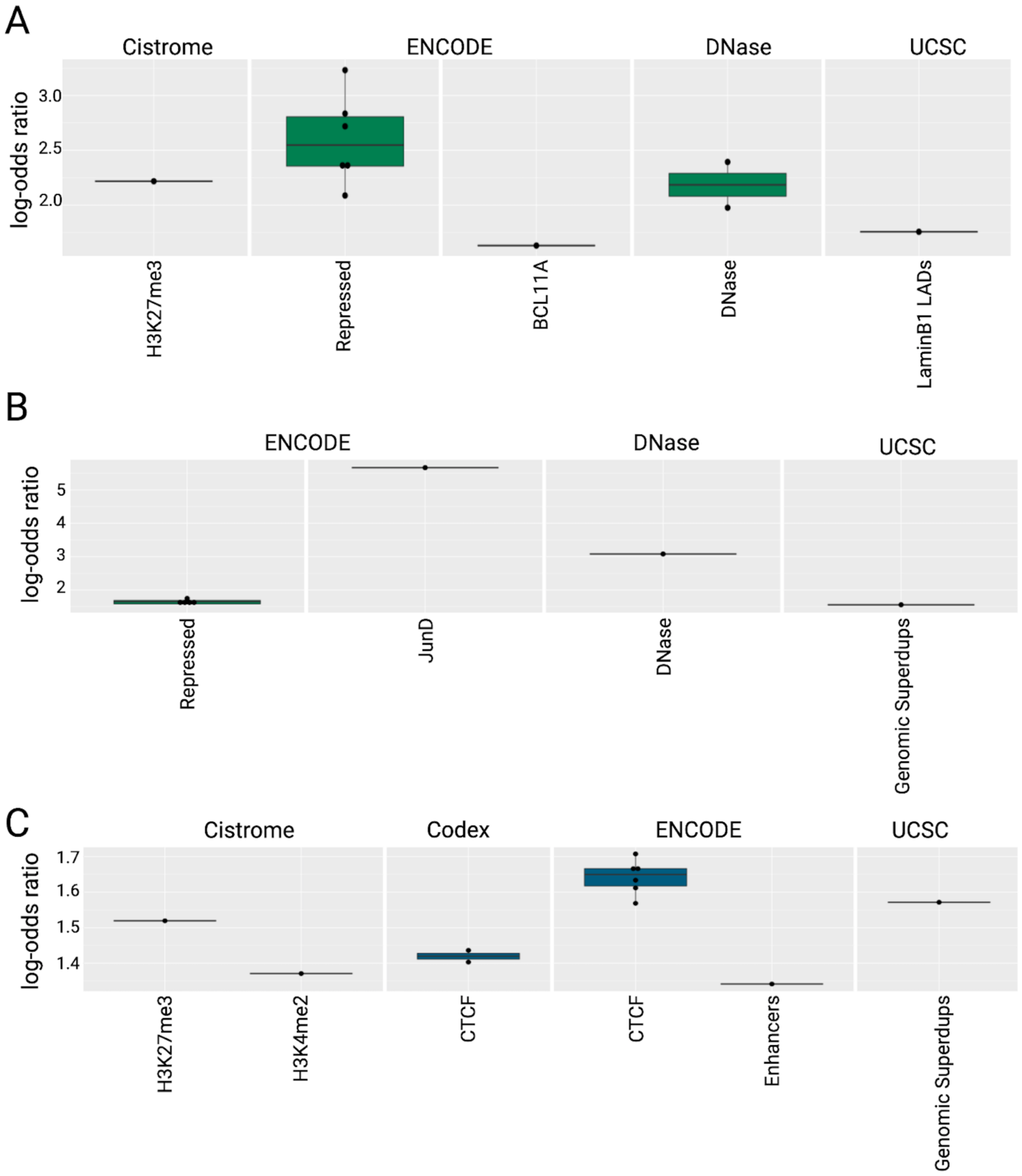


**Supplemental Figure S4. LOLA enrichment in hypermethylated regions in Peruvians with high vs. low methylmercury exposure.** Boxplots showing log-odds ratios ($p<0.01$) for the 1000 best ranking regions in LOLA enrichment analysis for (A) genomic tiling regions, (B) promoters, and (C) CpG islands that are hypermethylated in Peruvian study participants with high (>10 μg/g) vs. low (<1 μg/g) total hair mercury, a proxy for methylmercury exposure.

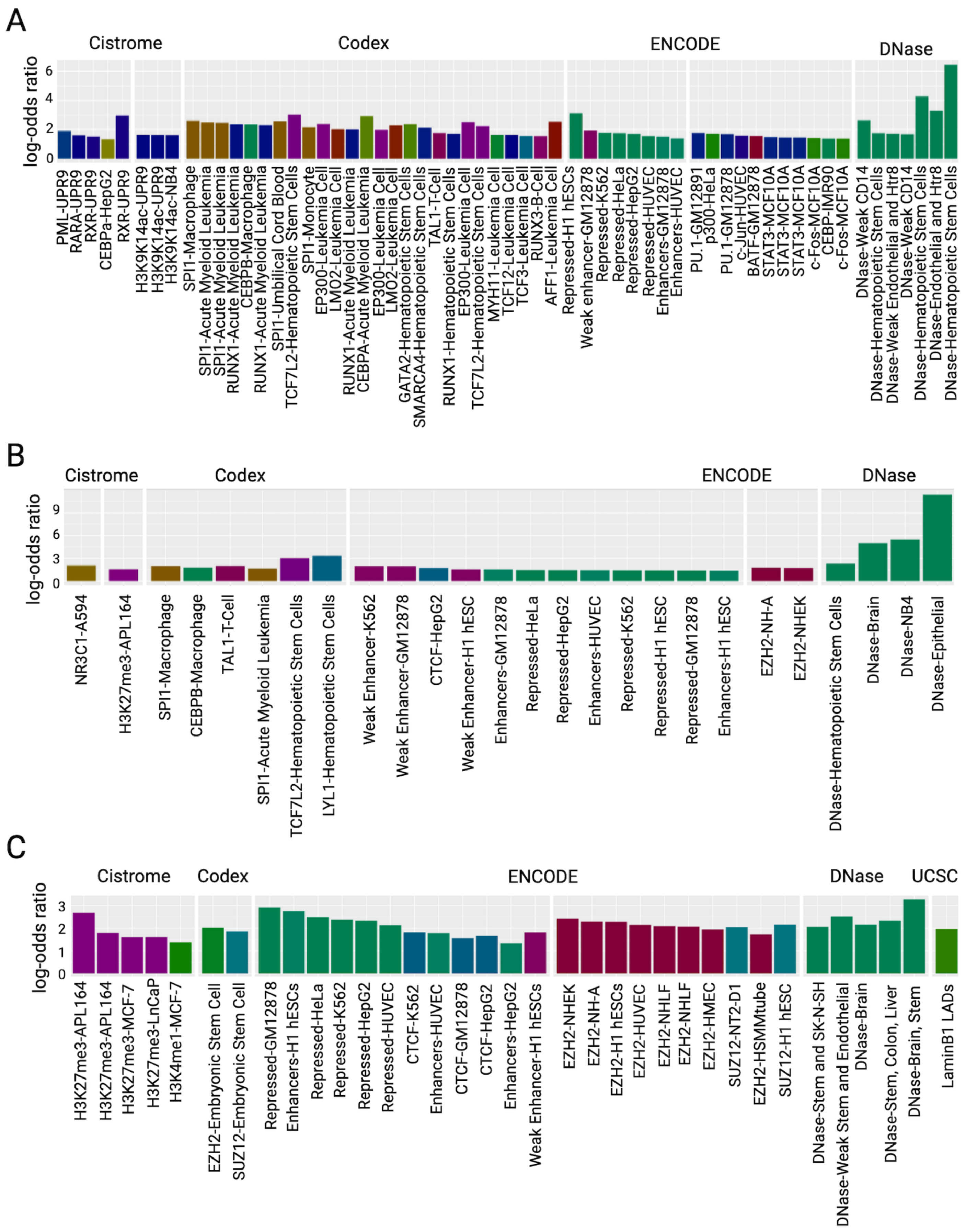


**Supplemental Figure S5. LOLA enrichment in hypomethylated regions in Peruvians with high vs. low methylmercury exposure.** Barplots showing log-odds ratios ($p<0.01$) for the 1000 best ranking regions in LOLA enrichment analysis for (A) genomic tiling regions, (B) promoters, and (C) CpG islands that are hypomethylated in Peruvian study participants with high (>10 $\mu$g/g) vs. low (<1 $\mu$g/g) total hair mercury, a proxy for methylmercury exposure.

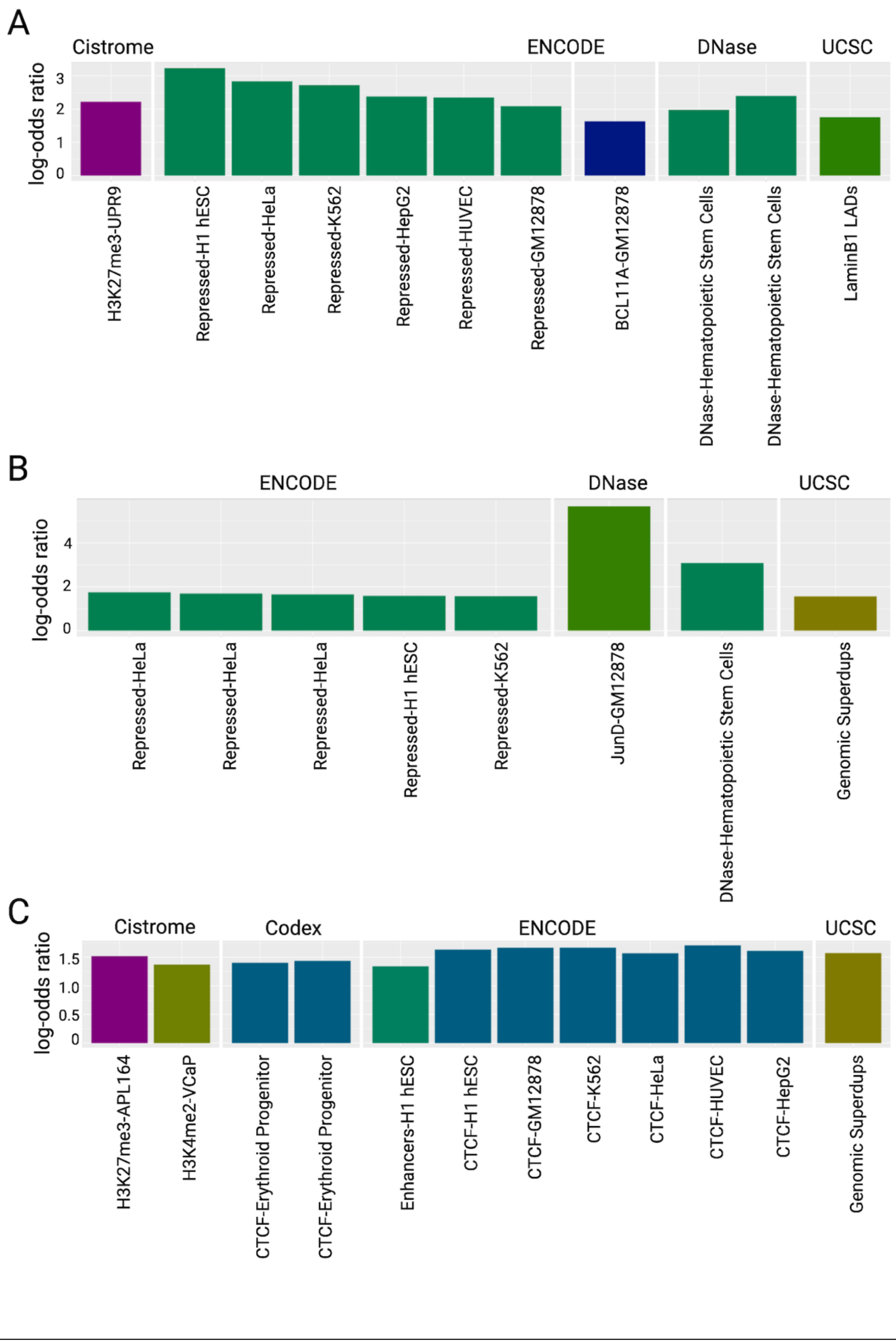


**Supplemental Figure S6. LOLA enrichment in hypermethylated regions in Peruvians with high vs. low methylmercury exposure.** Barplots showing log-odds ratios ($p<0.01$) for the 1000 best ranking regions in LOLA enrichment analysis for (A) genomic tiling regions, (B) promoters, and (C) CpG islands that are hypermethylated in Peruvian study participants with high (>10 μg/g) vs. low (<1 μg/g) total hair mercury, a proxy for methylmercury exposure.

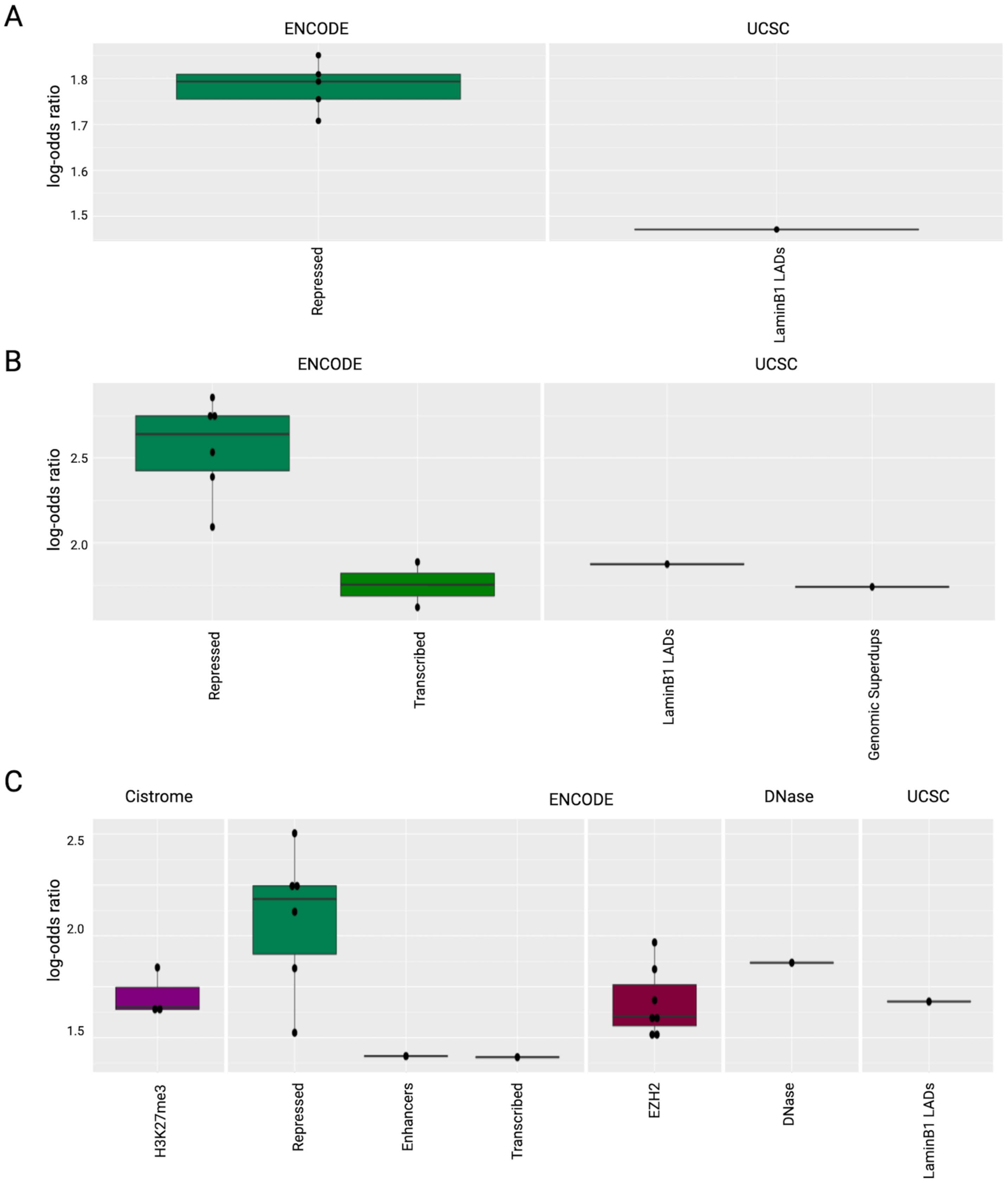


**Supplemental Figure S7. LOLA enrichment in regions with hypovariable DNA methylation in Peruvians with high vs. low methylmercury exposure.** Boxplots showing log-odds ratios ($p<0.01$) for the 1000 best ranking regions in LOLA enrichment analysis for (A) genomic tiling regions, (B) promoters, and (C) CpG islands that have hypovariable DNA methylation in Peruvian study participants with high (>10 μg/g) vs. low (<1 μg/g) total hair mercury, a proxy for methylmercury exposure.

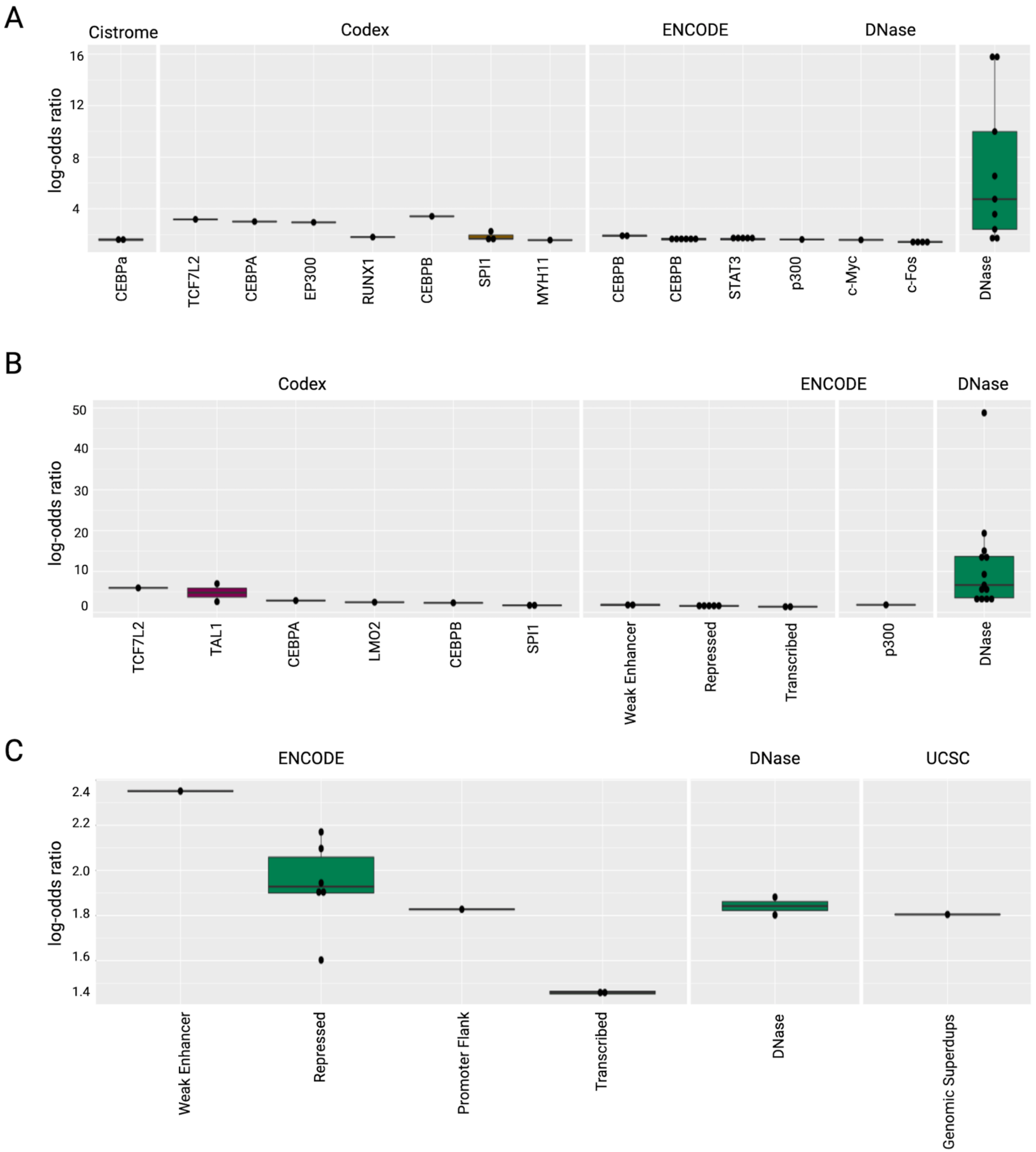


**Supplemental Figure S8. LOLA enrichment in regions with hypervariable DNA methylation in Peruvians with high vs. low methylmercury exposure.** Boxplots showing log-odds ratios ($p<0.01$) for the 1000 best ranking regions in LOLA enrichment analysis for (A) genomic tiling regions, (B) promoters, and (C) CpG islands that have hypervariable DNA methylation in Peruvian study participants with high (>10 μg/g) vs. low (<1 μg/g) total hair mercury, a proxy for methylmercury exposure.

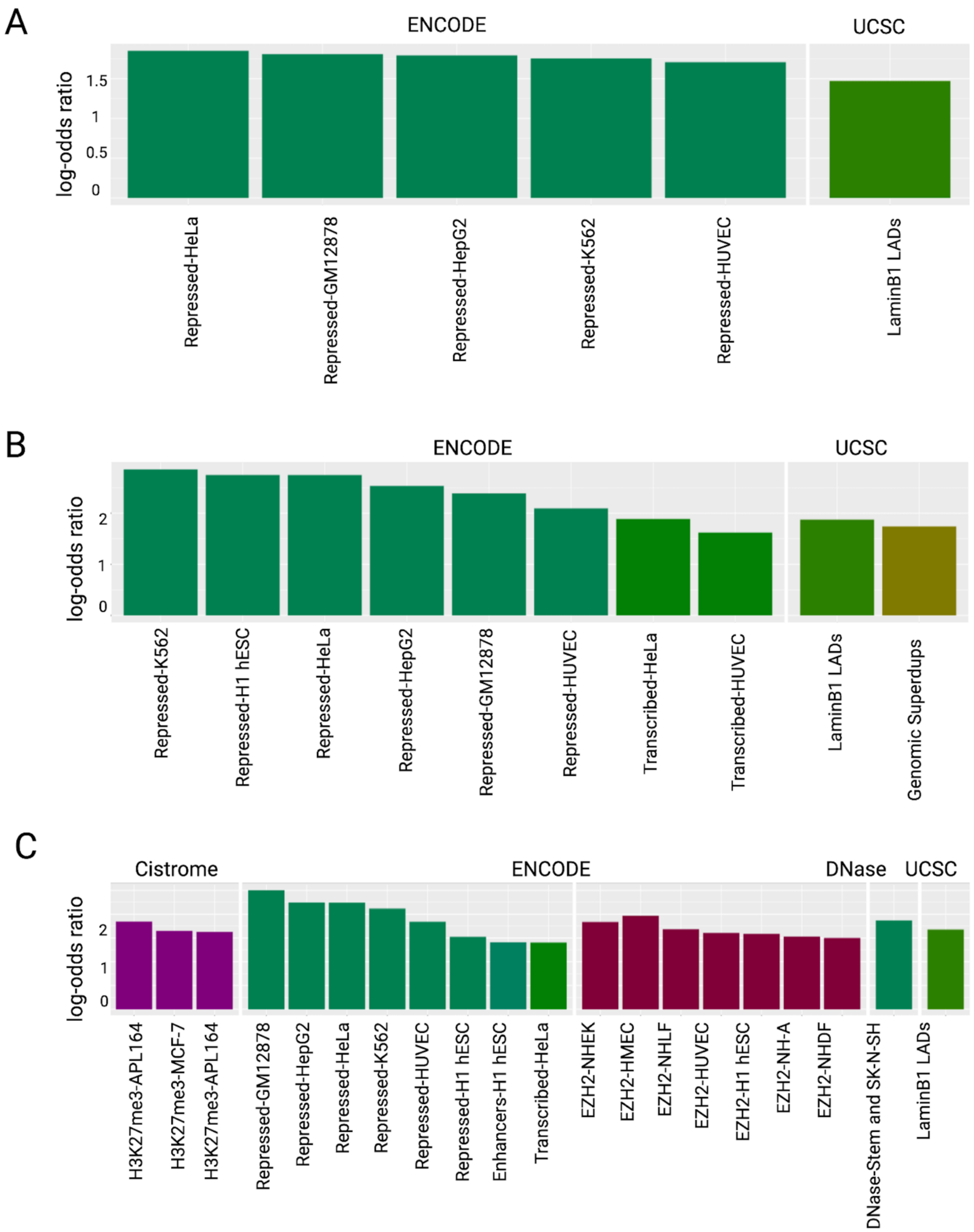


**Supplemental Figure S9. LOLA enrichment in regions with hypovariable DNA methylation in Peruvians with high vs. low methylmercury exposure.** Barplots showing log-odds ratios ($p<0.01$) for the 1000 best ranking regions in LOLA enrichment analysis for (A) genomic tiling regions, (B) promoters, and (C) CpG islands that have hypovariable DNA methylation in Peruvian study participants with high (>10 μg/g) vs. low (<1 μg/g) total hair mercury, a proxy for methylmercury exposure.

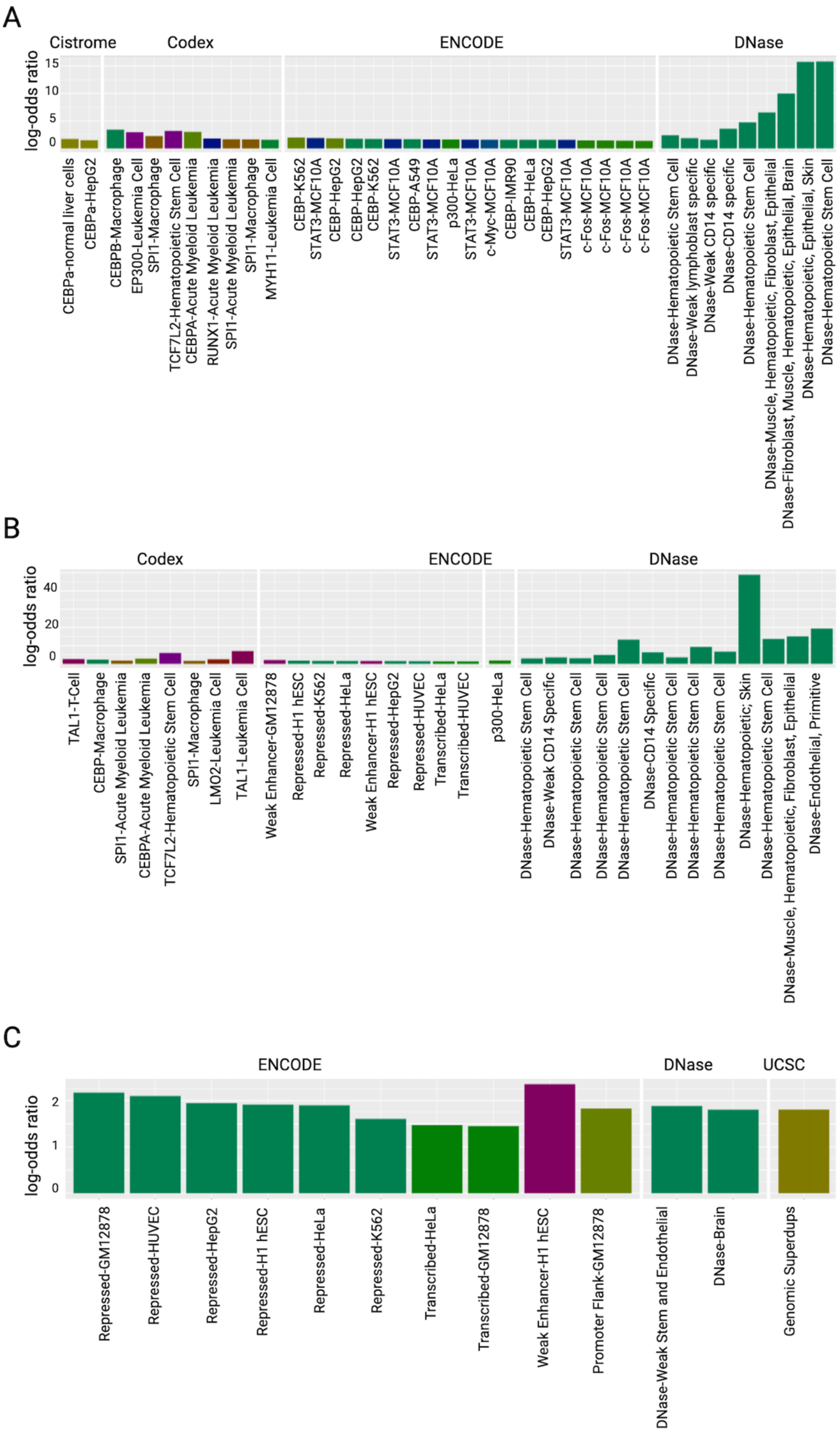


**Supplemental Figure S10. LOLA enrichment in regions with hypervariable DNA methylation in Peruvians with high vs. low methylmercury exposure.** Barplots showing log-odds ratios (p<0.01) for the 1000 best ranking regions in LOLA enrichment analysis for (A) genomic tiling regions, (B) promoters, and (C) CpG islands that have hypervariable DNA methylation in Peruvian study participants with high (>10 μg/g) vs. low (<1 μg/g) total hair mercury, a proxy for methylmercury exposure.

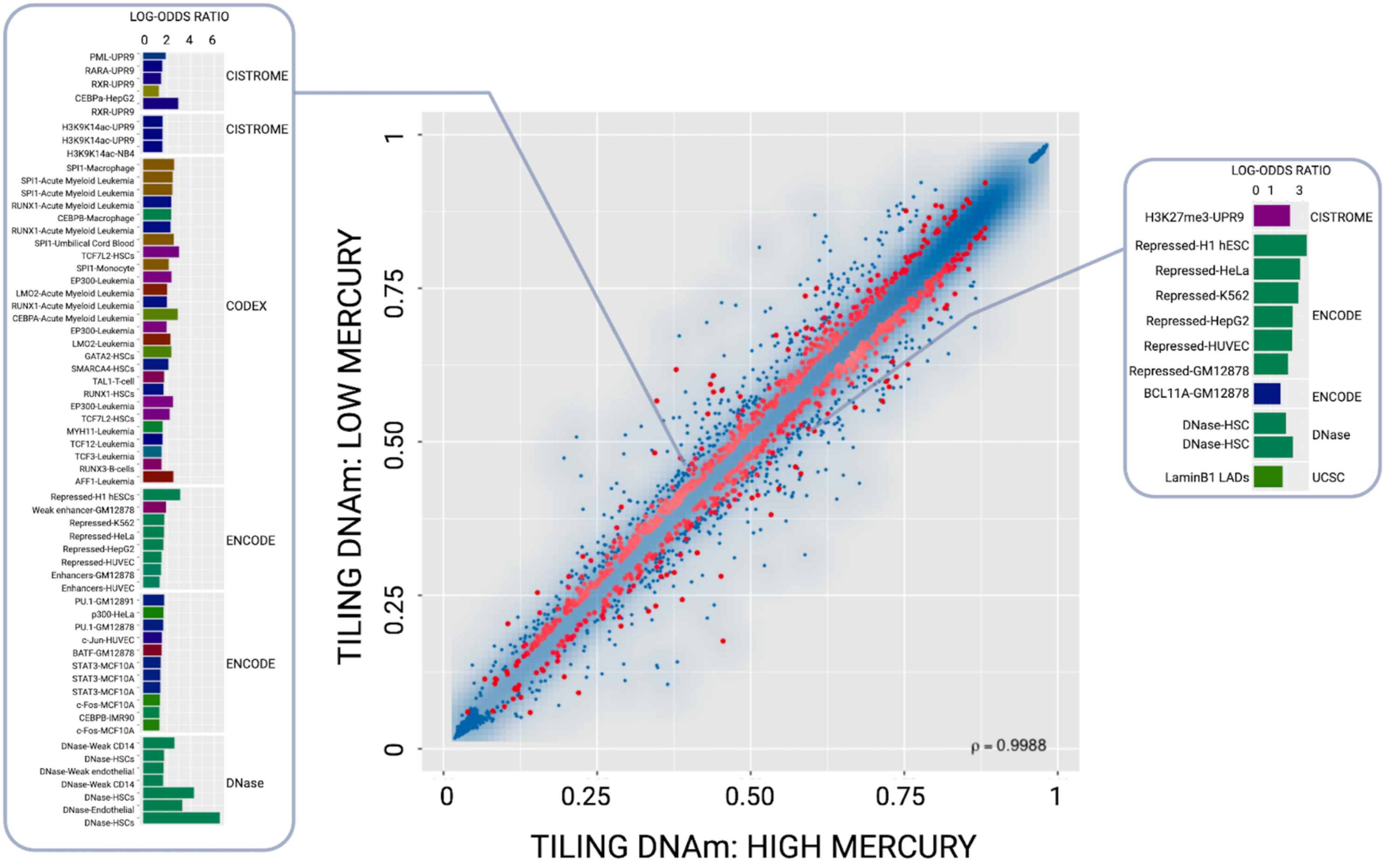


**Supplemental Figure S11. Transcription factor binding site enrichments in differentially methylated genomic tiling regions in individuals with high methylmercury exposure.** Scatterplot for differentially methylated genomic tiling regions. Color transparency corresponds to point density; the 1% of points in the sparsest population plot regions are drawn explicitly. Red colored points represent the 1000 best ranking regions. Barplots showing log-odds ratios ($p<0.01$) from LOLA enrichment analysis for genomic tiling regions that are hypomethylated (left) and hypermethylated (right) in Peruvian study participants with high (>10 μg/g) vs. low (<1 μg/g) total hair mercury, a proxy for methylmercury exposure. Barplots are additionally shown in Supplemental Figs. S5A and S6A.

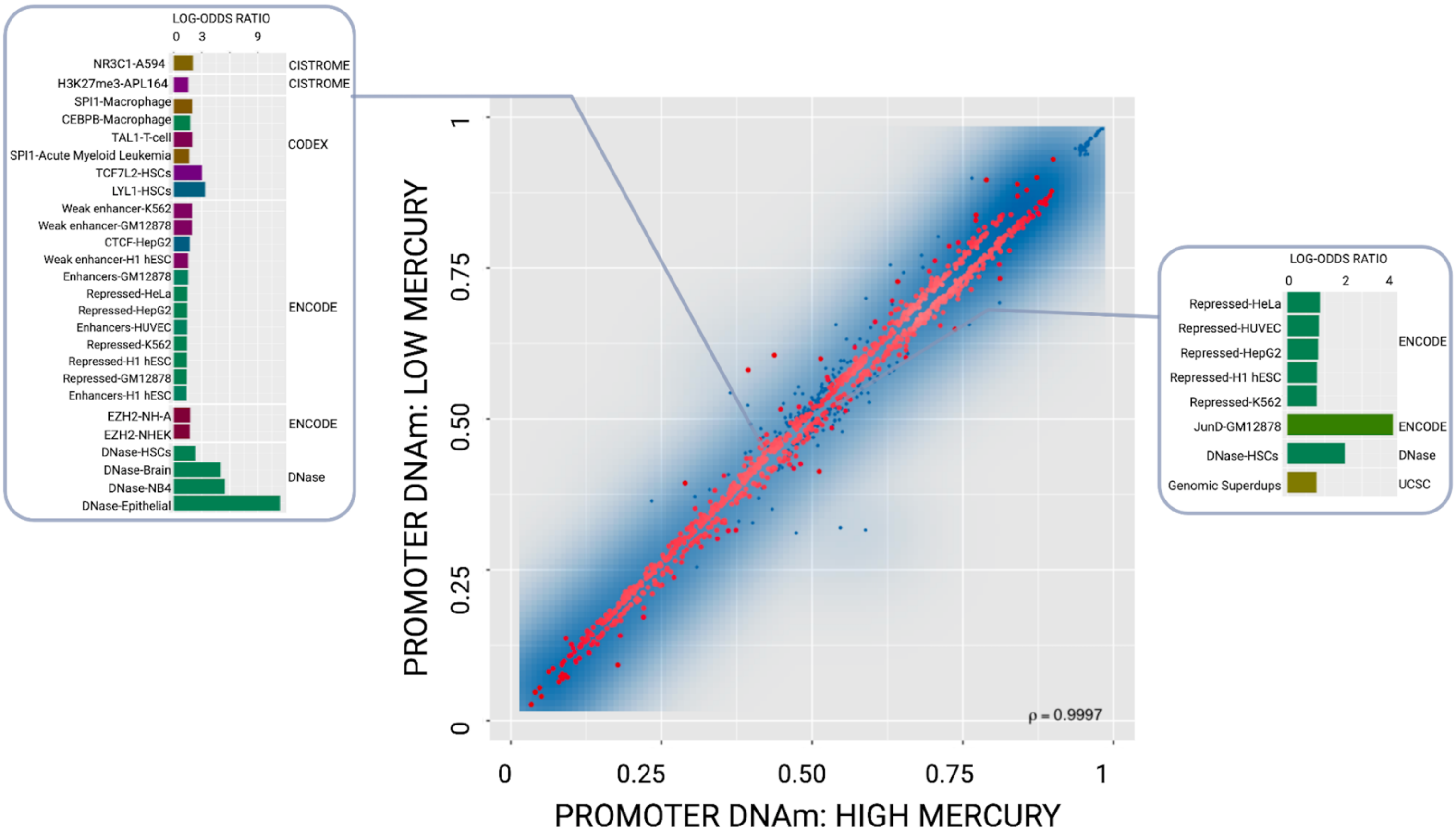


**Supplemental Figure S12. Transcription factor binding site enrichments in differentially methylated promoters in individuals with high methylmercury exposure.** Scatterplot for differentially methylated promoter regions. Color transparency corresponds to point density; the 1% of points in the sparsest population plot regions are drawn explicitly. Red colored points represent the 1000 best ranking regions. Barplots showing log-odds ratios ($p<0.01$) from LOLA enrichment analysis for promoter that are hypomethylated (left) and hypermethylated (right) in Peruvian study participants with high (>10 μg/g) vs. low (<1 μg/g) total hair mercury, a proxy for methylmercury exposure. Barplots are additionally shown in Supplemental Figs. S5B and S6B

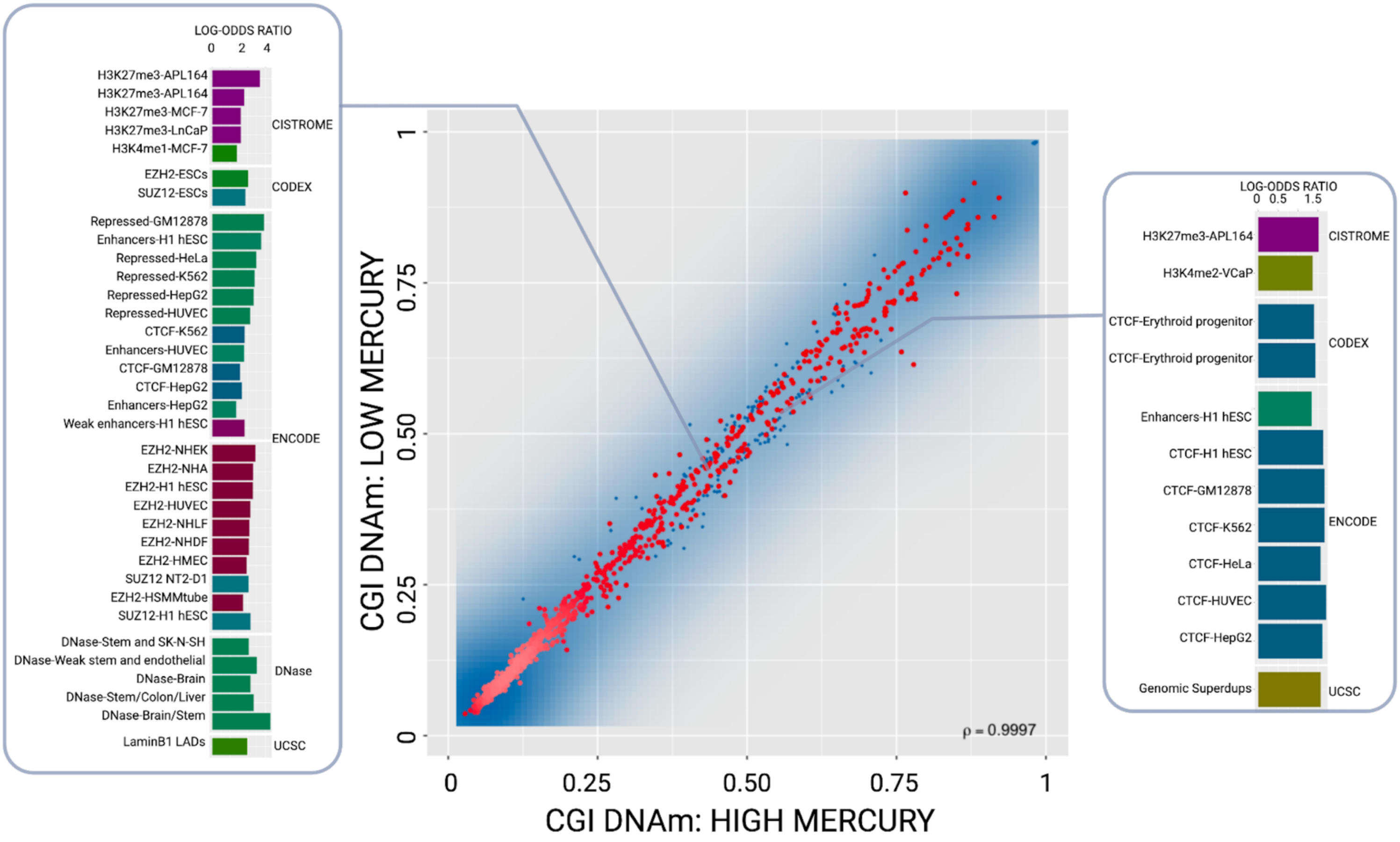


**Supplemental Figure S13. Transcription factor binding site enrichments in differentially methylated CpG islands in individuals with high methylmercury exposure.** Scatterplot for differentially methylated CpG island regions. Color transparency corresponds to point density; the 1% of points in the sparsest population plot regions are drawn explicitly. Red colored points represent the 1000 best ranking regions. Barplots showing log-odds ratios ($p<0.01$) from LOLA enrichment analysis for CpG islands that are hypomethylated (left) and hypermethylated (right) in Peruvian study participants with high (>10 μg/g) vs. low (<1 μg/g) total hair mercury, a proxy for methylmercury exposure. Barplots are additionally shown in Supplemental Figs. S5C and S6C.
